# dnoise: Fast Native Data Reduction for Bruker timsTOF

**DOI:** 10.64898/2026.08.27.747603

**Authors:** Patrick T. Garrett, Jolene K. Diedrich, John R. Yates

**Affiliations:** Integrated Computational, and Structural Biology, The Scripps Research, Institute, 10550 North, Torrey Pines Road, La Jolla, CA 92037, USA; Multi-Omics Core, The, Scripps Research, Institute, 10550 North, Torrey Pines Road, La Jolla, CA 92037, USA

**Keywords:** timsTOF, ion mobility, PASEF, denoising, proteomics, Rust

## Abstract

Bruker timsTOF acquisitions produce dense native `.d` files whose storage, transfer, and archival become substantial at high throughput. We present dnoise, an open-source Rust tool that removes points directly from timsTOF frames and writes a native-compatible `.d` directory. dnoise retains ions that form coherent streaks across the ion-mobility dimension and applies acquisition-aware gates to signal that cannot be selected for fragmentation. On a three-species benchmark spanning ddaPASEF and diaPASEF at 5- and 15-minute gradients, default MS1-only denoising reduced the frame binary by 35 to 53%. Label-free quantification accuracy was preserved in both modes. ddaPASEF peptide-spectrum-match, peptide, and protein-group counts were unchanged, as expected with the searched MS/MS spectra untouched, and diaPASEF precursor and protein-group counts changed only slightly. Every tested processing run completed in 69 seconds or less on the benchmark workstation. Optional MS/MS denoising produced greater reduction but sacrificed several percent of identifications. Thus, a substantial fraction of native timsTOF frame data can be removed with little analytical change.

dnoise at a glance, on real data: a representative MS1 frame from a 5-minute ddaPASEF run, with retained points in teal and discarded points in grey. For this display the precursor-selection gate is applied with no edge padding (the literal selection polygon), where the benchmark configuration pads its edges to protect the isotopic envelopes of edge precursors. The headline outcomes summarize default MS1-only denoising across ddaPASEF and diaPASEF at both gradients.

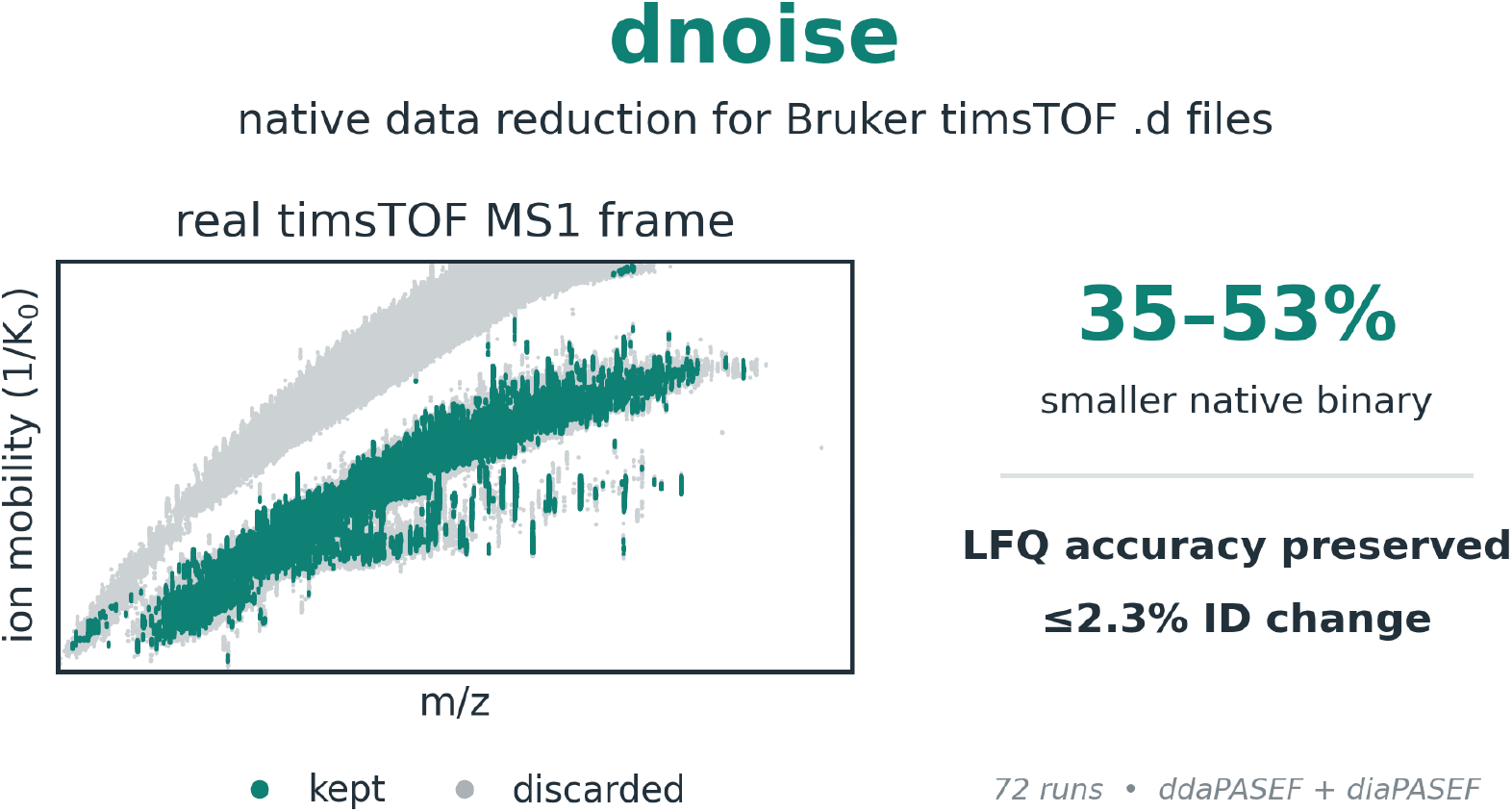

## 1 Introduction

Bruker timsTOF instruments combine trapped ion mobility spectrometry (TIMS)^1,2^ with parallel accumulation–serial fragmentation (PASEF)^3^ to record retention time, mass-to-charge 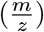, ion mobility, and intensity of detected ions. These measurements are stored in native `.d` directories as thousands of frames, each containing a complete mobility ramp and often hundreds of thousands of digitized points. The resulting data volume is an operational cost for high-throughput proteomics. The densest files in this study contained 6.7 GB of frame data over 27 minutes of acquisition, including loading and washout around the 15-minute gradient. Running that method continuously would produce about 0.36 TB per day or 130 TB per year, before redundant backups and public deposition.

A substantial fraction of these points do not contribute to the intended analysis. Both data-dependent acquisition (DDA, ddaPASEF) and data-independent acquisition (DIA, diaPASEF) define a fragmentable region in the 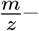 mobility plane: a precursor-selection polygon in ddaPASEF and isolation windows in diaPASEF. These boundaries exclude the singly charged ion cloud and regions where multiply charged peptides are not expected, but they govern fragmentation rather than recording. MS1 frames therefore retain signal that can never be selected. In representative 5-minute runs, 50.1% of ddaPASEF and 72.7% of diaPASEF MS1 points lay outside the fragmentable region.

Recorded points also differ in mobility structure. Peptide ions appear as dense streaks at nearly constant 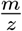 across consecutive mobility scans, whereas sparse points and diffuse clusters cannot be resolved to a useful feature. A second artifact forms a halo of weak signal around intense peaks, consistent with microchannel-plate saturation in time-of-flight instruments^4^. These patterns provide a structural basis for removing points without applying an absolute per-point intensity cutoff.

Existing approaches do not directly fill this role. Peak detection and feature finding commonly operate after conversion to mzML^5,6^, producing an intermediate representation rather than a smaller native file. Numerical and general-purpose compression re-encode the data but preserve the uninformative points, while the frame binary is already compressed. On other ion-mobility platforms, the PNNL PreProcessor writes denoised data back in the instrument’s own format^7^, but it does not support Bruker `.d`. Search pipelines, access libraries, and processing frameworks read Bruker `.d` data natively^8–13^, but each emits search results or its own representation of the data rather than a reduced `.d`. The spectral-simplification method of Wilding-McBride et al.^14^ exports MGF or feature lists. To our knowledge, no open-source tool removes points and writes the result as a new native-compatible Bruker `.d` directory.

Here we present dnoise, an open-source Rust tool that combines an ion-mobility streak filter, removal of weak peak halos, and acquisition-aware geometric gates. Frames are processed independently and written to a new `.d` directory in the native format, leaving the original untouched. We evaluate the default MS1-only mode on a defined human/yeast/*E. coli* mixture across 5-and 15-minute gradients in ddaPASEF and diaPASEF. It reduced the native frame binary by 35–53% while preserving LFQ accuracy in both modes. ddaPASEF peptide-spectrum match (PSM), peptide, and protein-group counts were unchanged, and diaPASEF precursor and protein-group counts changed only slightly. We also characterize optional MS/MS filtering as a higher-reduction operating point with an identification tradeoff.

## 2 Experimental Section

### 2.1 dnoise filtering

dnoise represents each frame as integer triples (*s, t, I*): mobility-scan index, time-of-flight (TOF) index, and intensity. Ion mobility and 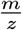 are recovered from the run calibration in `analysis.tdf`, but the filters operate in stored index space. Calibration is used only to translate acquisition regions specified in physical units into scan and TOF-index boundaries.

For each TOF column, the streak filter sums intensities over a neighboring TOF window to form a per-scan mobility profile. Occupied scans are grouped into runs, bridging gaps up to `max_internal_gap`. A run is retained when it contains at least `min_feature_length` occupied scans and satisfies the optional intensity floors. Points in retained runs survive, and the procedure repeats for `iterations` passes. At the defaults used here, both intensity floors are zero, so the decision is based on mobility occupancy and length rather than absolute intensity. Complete defaults are listed in Table S1.

The halo filter acts on the streak-filter survivors. For each point it finds the maximum intensity in a local box while excluding the point’s own TOF column, which protects the ion’s mobility streak. A point is removed when its intensity is below `halo_peak_fraction` (0.15) of that off-column maximum. The filter can be disabled because a sufficiently weak co-eluting ion could meet the same criteria.

Acquisition-aware gates remove MS1 points outside the fragmentable region stored in `analysis.tdf`: the precursor-selection polygon for ddaPASEF and the isolation-window table for diaPASEF. Both accept padding in 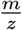 and mobility to protect edge features. Matching gates remove fragment points outside their isolation events when optional MS/MS filtering is enabled. Gates silently do nothing when their defining method table is absent, allowing the same default configuration to select the appropriate behavior for either acquisition mode. Figure 1 shows the three default stages applied to a representative MS1 frame.

**Figure 1:**
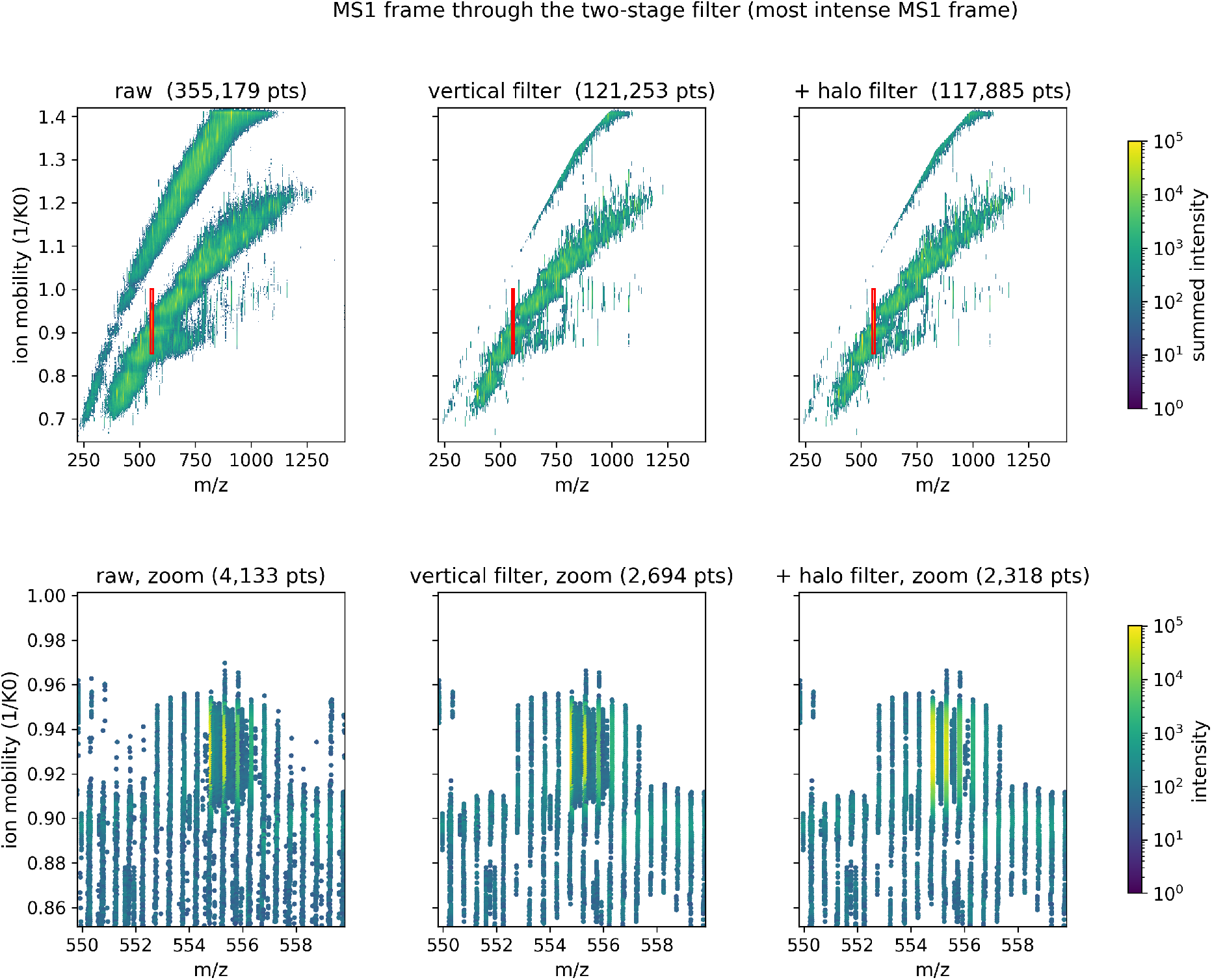
The most intense MS1 frame from a representative 5-minute ddaPASEF run through the default filter stages. Columns show the raw frame (355,179 points), the acquisition gate plus ion-mobility streak filter (121,253), and the added 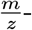 halo filter (117,885). The top row shows the full 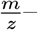 mobility frame. The bottom row enlarges the outlined region.

### 2.2 Parameter selection

The compiled defaults in Table S1 were held fixed across the four tested acquisition-mode and gradient combinations. The two geometric parameters of the streak filter, `min_feature_length` and `max_internal_gap`, were selected by a grid sweep on one homogeneous sample, the six replicates of Condition A of the 15-minute ddaPASEF gradient, scored on quantified coverage, replicate precision, and intensity fidelity to the unfiltered data (Table S2). The chosen setting removed a large fraction of MS1 points while quantifying slightly more peptides and proteins than the unfiltered data at unchanged precision. The sweep was a selection aid rather than an objective to maximize benchmark coverage. We chose gap 2 and length 5 to prioritize stricter local continuity and greater point removal over maximizing coverage on the selection sample. Both intensity floors were left at zero. Because the selection sample is part of the benchmark, the 15-minute ddaPASEF results are not fully out-of-sample. The 5-minute gradient and both diaPASEF acquisitions played no role in parameter selection.

### 2.3 Optional MS/MS filtering

The default mode filters MS1 survey frames only. An optional stage applies the streak and halo filters within each fragment isolation event, using relaxed `msms_*` parameters because fragment spectra are sparser. It is disabled by default because removing fragment peaks can change identifications.

For ddaPASEF, dnoise combines scans from every frame that re-isolated a precursor, filters the summed spectrum, and maps the surviving scan/TOF coordinates back to each native frame. For diaPASEF, which has no corresponding precursor table, each isolation window is filtered independently. This prevents mobility runs from joining across windows representing different precursor populations. Representative frames and parameter sweeps are in Section S4.

### 2.4 Implementation and native-format compatibility

dnoise is implemented in Rust as a command-line tool and library. It reads frames with `timsrust` 0.4.2^15^, filters and encodes them in parallel with `rayon`, and writes the Bruker type-2 `analysis.tdf_bin` encoding with its own encoder. For each frame it also updates the byte offset, peak count, maximum intensity, and summed intensity recorded in the accompanying `analysis.tdf` database. An unfiltered read/write round trip through dnoise was point-for-point identical when decoded with `timsrust`. The default filters are equally conservative, retaining or discarding native points without moving them or changing their intensities.

### 2.5 Benchmark design

We used the Generation Beta three-species hybrid benchmark (human, *Saccharomyces cerevisiae*, and *Escherichia coli*) deposited as PRIDE PXD070049^16–19^. Its defined ratios provide a standard test of label- free quantification accuracy^20^. Conditions A, B, and C contain human/yeast/*E. coli* at 65/30/5%, 65/15/20%, and 65/3/32%, producing expected pairwise log_2_ ratios from −2.7 to +3.3 (Table S3). We analyzed timsTOF Ultra 2 ddaPASEF and diaPASEF acquisitions at 5- and 15-minute gradients with a 50 ng load. Each condition had six replicates, for 18 runs per gradient and acquisition mode (72 raw runs total).

### 2.6 Database search and quantification

The original, MS1-only, and MS1+MS/MS ddaPASEF arms were searched with Sage 0.15.0-beta.1^21^ and the same species-tagged FASTA and configuration. Searches were fully tryptic, allowed two missed cleavages and peptide lengths of 7–30, used fixed cysteine carbamidomethylation and variable methionine oxidation, and applied ±20 ppm precursor and fragment tolerances. Sage provided mobility-aware MS1 label-free quantification: precursor signals were integrated across retention time and mobility, including charge states and isotopologues, and identifications were transferred between runs using decoy-controlled LFQ q-values, as in IonQuant^22^.

The three diaPASEF arms were searched independently with DIA-NN 2.2.0^23^, which read each native `.d` directory through its bundled Bruker `timsdata` library. All arms used one whole-proteome spectral library predicted from the benchmark FASTA before denoising. Each arm built its own two-pass refinement and report. DIA-NN quantified proteins with MaxLFQ^24^ from MS2 fragment chromatograms. A direct MS1 check used its `Ms1.Normalised` values, summed per precursor and run.

In both modes, quantified proteins required at least two distinct quantified peptides and measurements in at least two replicates of each compared condition. Each run was normalized to the cross-run mean total intensity within its gradient, consistent with equal total protein loading. Complete configurations and software versions are provided in Section S7.

### 2.7 Evaluation metrics

We measured reduction of the native `analysis.tdf_bin`, retention of PSMs or precursors and protein groups, LFQ accuracy, runtime, and memory. We also report quantified protein and peptide counts and within-condition precision. Accuracy is the species-specific median log_2_ ratio relative to the known mixture ratio. For paired original-versus-MS1 accuracy comparisons, percentile-bootstrap 95% confidence intervals were calculated by resampling shared contributing proteins (2,000 resamples). The corresponding intervals are reported in Tables S10 and S11. Precision is the median pooled protein CV across conditions A and B.

## 3 Results and Discussion

### 3.1 dnoise substantially reduces timsTOF data volume

At its default settings (Table S1), dnoise removed the large majority of MS1 points in every acquisition tested. The effect was consistent across all 18 runs of each condition and across the 5- and 15-minute gradients (complete results in Table S5 and Section S2). In ddaPASEF at 5 and 15 minutes, respectively, MS1 peaks fell by 81.2% and 78.2%, and the native `analysis.tdf_bin` frame binary fell by 53.4% and 49.5% (Figure 2). The 15-minute runs contained 2.1 times as many raw MS1 points as the 5-minute runs, yet the fraction removed changed little.

**Figure 2:**
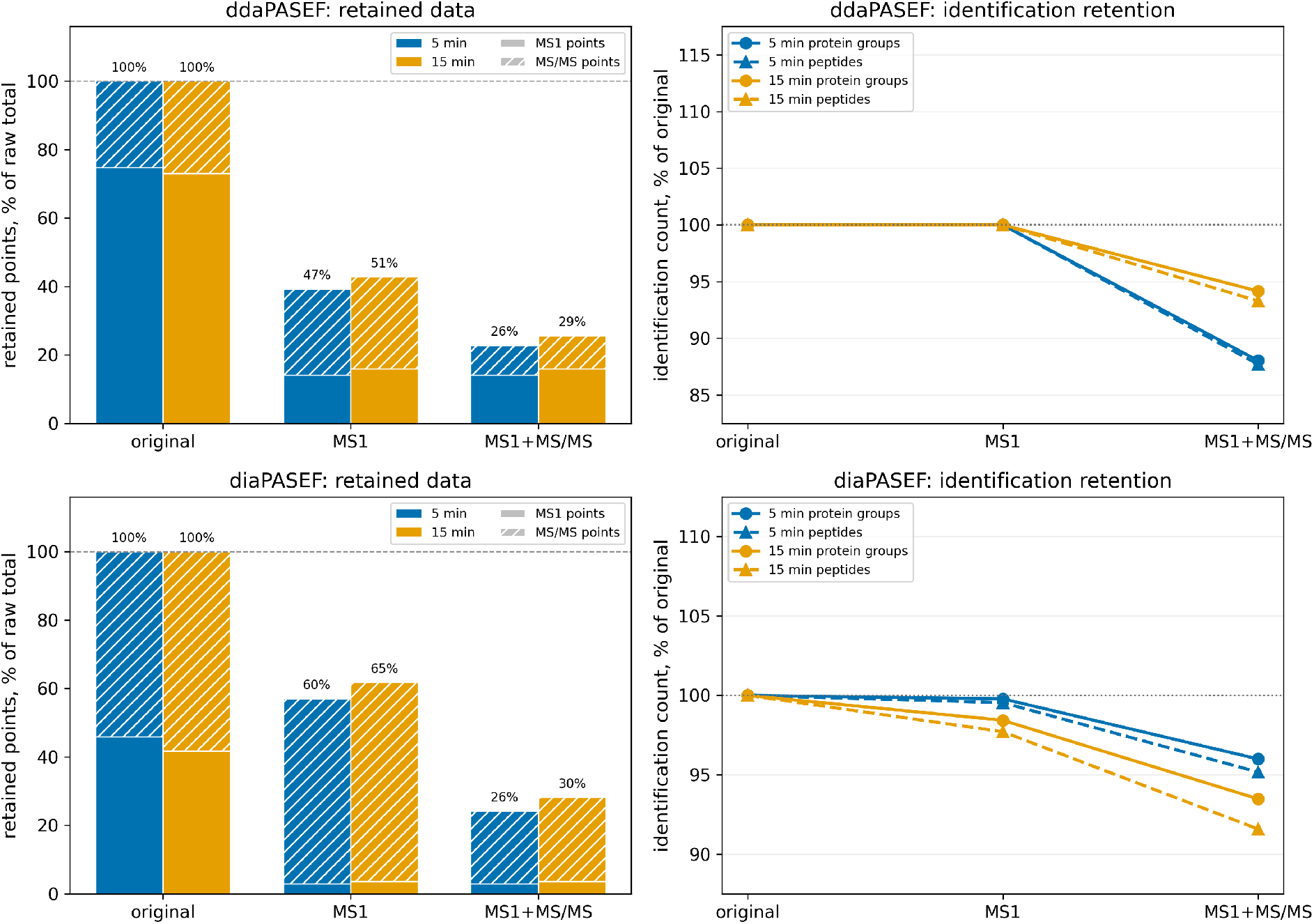
Data reduction and identification retention across acquisition modes. **Left:** retained points partitioned into MS1 and MS/MS contributions, with bar-top labels giving the retained frame-binary size. **Right:** identified protein-group and peptide counts relative to the original arm, for the default MS1-only and optional MS1+MS/MS arms. Complete values are in Table S5 and Table S6.

The reduction was larger in diaPASEF. At 5 and 15 minutes, respectively, MS1 denoising removed 93.5% and 91.4% of MS1 points, and the frame binary fell by 39.6% and 34.7% (Figure 2). This difference should not be interpreted as an inherent advantage of one acquisition mode. Bruker’s on-instrument denoising was enabled for the ddaPASEF survey scans but not for the diaPASEF scans, so dnoise encountered more background in the latter. Within each acquisition, reduction was reproducible across runs and gradient lengths. A per-stage ablation on one run per acquisition separates the three default components (Table S4). The acquisition gate and the streak filter each remove a large fraction of MS1 points on their own, and their combination removes most of the total. The halo filter is a small final trim.

### 3.2 Default MS1 denoising preserves the analytical result

In ddaPASEF, the original and MS1-denoised arms returned identical PSM, peptide, and protein-group counts at both gradients (Figure 2, Table S5), as expected, since the database search reads only the MS/MS spectra, which the default mode leaves untouched. The substantive ddaPASEF validation is therefore quantitative, and label-free quantification was preserved. Across all three species and all condition pairs, the original and MS1-denoised arms followed the same expected ratios over the full benchmark range (Figure 3). The number of quantified proteins and median within-condition CV changed slightly, but neither accuracy nor precision showed a systematic degradation attributable to denoising. At 5 minutes, the largest movements for the regulated species were toward their expected ratios. The smaller changes across the remaining ddaPASEF cells and diaPASEF were mixed. Complete identification, quantification, accuracy, and precision values are reported in Section S3.

**Figure 3:**
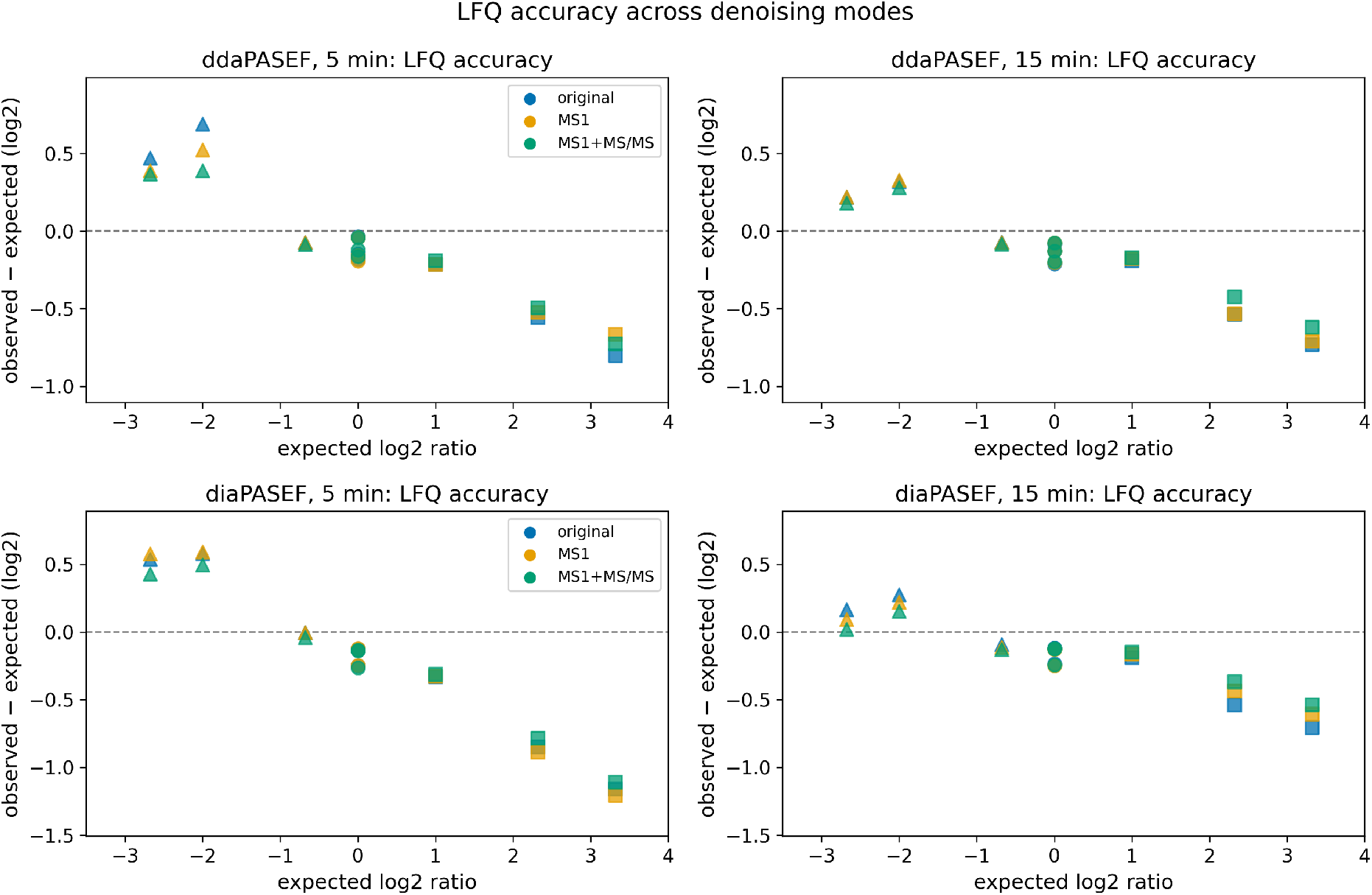
LFQ accuracy across acquisition modes. Each panel shows residual median log_2_ ratios (observed minus expected) for all condition pairs and species, where zero is ideal. Precision for the same arms is in Figure 4, and complete values are in Section S3.

MS1 denoising had a small identification cost in diaPASEF. At 5 and 15 minutes, respectively, experiment-wide precursor counts changed by 0.2% and 2.2%, and protein-group counts changed by 0.2% and 1.6% (Figure 2). Median protein-level LFQ CV and LFQ accuracy (Figures 3 and 4) were unchanged at both gradients. DIA-NN derives the reported MaxLFQ quantity from fragment chromatograms, so the default MS1-only filter does not directly alter the signal used for quantification. A separate check of the direct precursor MS1-area field showed that the regulated-species ratios moved toward their expected values after denoising, with only modest changes in precursor CV (Table S12). The file therefore becomes smaller while the fragment-based protein ratios remain stable and the direct precursor-area ratios move toward expectation.

**Figure 4:**
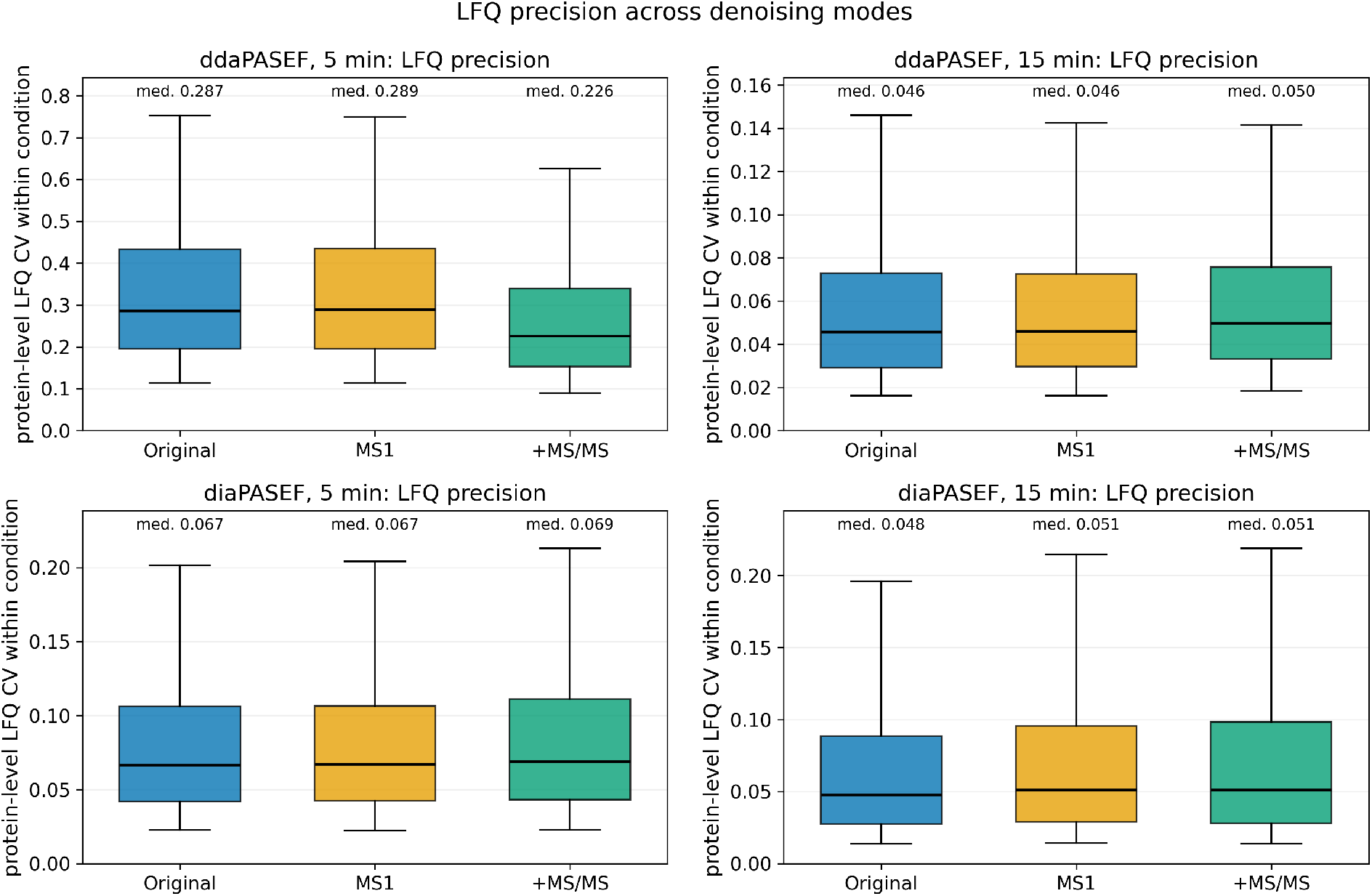
LFQ precision across acquisition modes. Each panel shows protein-level CV distributions within conditions A and B. Whiskers span the 5th–95th percentiles. Complete values are in Section S3.

Precision was likewise preserved under default MS1 denoising. Protein-level CV distributions and their medians changed only slightly at either gradient or acquisition mode (Figure 4).

Feature-level intensities provided a more direct fidelity check than summary ratios alone. Across both gradients and all replicate runs, shared DDA peptide and DIA precursor intensities closely followed the equality line after default MS1 denoising (Figure 5). Agreement remained high with optional MS/MS denoising.

**Figure 5:**
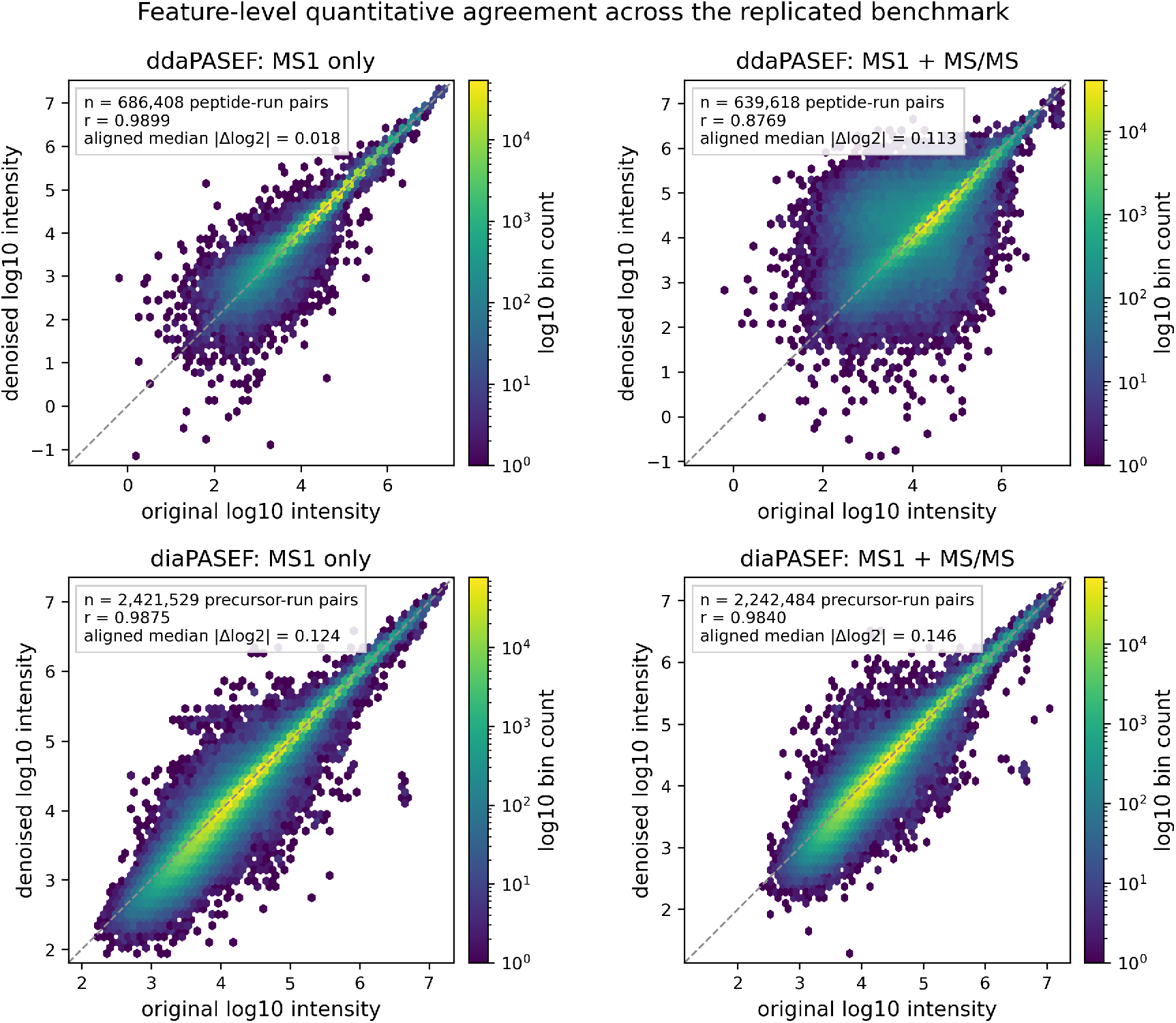
Feature-level quantitative agreement across the replicated benchmark. Each hexagon pools positive, original–denoised feature/run pairs from the 5- and 15-minute gradients. Pearson *r* is calculated on log_10_ intensities. The aligned median absolute Δ log_2_ is calculated after centering paired log-ratio distributions on their medians.

### 3.3 Optional MS/MS denoising trades identifications for greater reduction

The default configuration filters MS1 frames only. dnoise can also filter MS/MS frames when storage reduction is more important than complete identification retention. In ddaPASEF at 5 and 15 minutes, respectively, this reduced the frame binary by 74.0% and 70.7%, while identified peptide counts fell by 12% and 7% (Figure 2). In diaPASEF at 5 and 15 minutes, respectively, the same option reduced the frame binary by 74.3% and 69.7%, with identified precursor and protein-group losses of several percent (Figure 2, Table S6).

Identification and quantification moved differently under this mode, and Table S5 contains an apparent inconsistency worth explaining: at the 5-minute gradient the MS1+MS/MS arm quantified more proteins and peptides than the original arm, at lower median CV, despite identifying fewer peptides. The increase reflects quantification eligibility, not recovered signal. A peptide enters the quantified set through Sage’s decoy-controlled LFQ q-value, which gates the transfer of identifications between runs. Fragment filtering removes weakly supported identifications first (they are enriched for false matches, Figure S6), and those are also the peptides that fail cross-run transfer. The surviving candidate pool is smaller but better behaved: the fraction passing the 1% LFQ q-value rose from 54% to 65%, and the peptides that passed were measured in more runs, so more proteins cleared the two-peptide, two-replicate reporting rule. The smaller quantified-count movements in the MS1-only arms (Table S5, Table S7) follow the same logic.

This operating point is useful but not analytically neutral. Fragment peaks provide the evidence for peptide identification, so removing them inevitably trades sensitivity for size. We therefore recommend MS1-only denoising as the default and present MS1+MS/MS denoising as an optional, tunable mode for users who can accept identification loss. Parameter sweeps supporting this tradeoff are provided in Section S4.

### 3.4 The streak filter better preserves weak signal than a matched intensity threshold

As a control, we compared the streak filter with a strict per-point intensity threshold calibrated to remove approximately the same fraction of points. Both arms applied the same acquisition-aware gates, so they differed only in the filter itself. The threshold retained a point only when its own intensity cleared a cutoff, using no information from neighboring mobility scans. For this removal-matched control, we calibrated that cutoff separately for each tested acquisition while holding the streak-filter configuration fixed. At comparable removal the two approaches produced similar MS1 reduction and broadly similar summary results when only MS1 frames were filtered.

Within shared ddaPASEF peptide-run pairs at both gradients, the streak filter increasingly outperformed the per-point threshold as abundance fell. Streak-filtered LFQ intensities stayed close to the original, whereas the threshold produced a progressively larger downward bias toward the faintest peptides (Figure S5). This abundance-dependent divergence is consistent with the streak filter retaining weak but mobility-coherent signal that a fixed per-point threshold discards.

Extended to fragment frames, the threshold also lost more quantified coverage than the streak filter, most clearly in diaPASEF (Figure S4, Section S5). The target-decoy search explains why. A rank-1 decoy hit is a match to a sequence known to be absent from the sample, an observed false positive rather than an estimated one. In the original ddaPASEF searches at 5 and 15 minutes, respectively, 30.1% and 23.7% of scored spectra had a decoy as their best available explanation. These spectra were never identifiable. MS1-only denoising reproduced the original search result exactly, but filtering the fragment frames removed the unidentifiable pool faster than the evidence. At 5 and 15 minutes, respectively, rank-1 decoy hits fell by 48.1% and 42.7%, while target hits fell by 27.2% and 18.2%, a decoy-to-target loss ratio of 1.77 and 2.34 (Figure S6). The matched intensity threshold inverted this ratio when extended to fragment frames, at 0.43 and 0.57: it lost real identifications faster than false ones. The identification loss under MS/MS denoising is therefore not a uniform tax: the spectra the streak filter withdraws are enriched for matches that were already wrong. Thus, the streak filter preferentially retained searchable signal, and the matched intensity filter did not.

### 3.5 Optional MS1 centroiding trades mobility resolution for extreme reduction

As an optional final stage, dnoise can replace groups of surviving MS1 points with intensity weighted centroids at newly calculated integer (scan, TOF-index) coordinates. The watershed centroider grows groups transitively and collapses each ion streak toward a single centroid, giving up the mobility profile for the largest reduction. The box centroider instead tiles each streak into small fixed boxes whose centroids preserve that profile at a more moderate reduction. On the 5-minute ddaPASEF benchmark, the watershed reduced MS1 to 2% of the raw points and the frame binary to 34% of raw. The box centroider reduced MS1 to 10% (binary 40%).

Despite collapsing MS1 to a small fraction of its points, both settings preserved the label-free result. Identifications were again identical, and precision improved slightly under both. The box centroider raised quantified coverage, while the watershed gave up 1.3% of the proteins and 2.2% of the peptides quantified by the original arm. Accuracy moved by at most 0.02 log_2_ units in the human and yeast ratios. The only larger movement was a small further de-compression of the low-abundance *E. coli* ratio. Both centroiders are off by default. They are high-reduction operating points, most useful when MS1 is needed only for precursor-level quantification at reduced mobility resolution. Both are characterized in full in the Supporting Information (Section S6, Figure S7, Table S15).

### 3.6 dnoise is fast enough for routine post-acquisition use

All processing times were shorter than even the shortest tested gradient of 5 minutes (Figure 6, Table S16). Across the 72 files, default MS1-only processing took 7.4–39.0 seconds on a single workstation (Intel Core i7-12700H, 20 threads). Processing MS1 and MS/MS frames took 10.2– 68.7 seconds.

**Figure 6:**
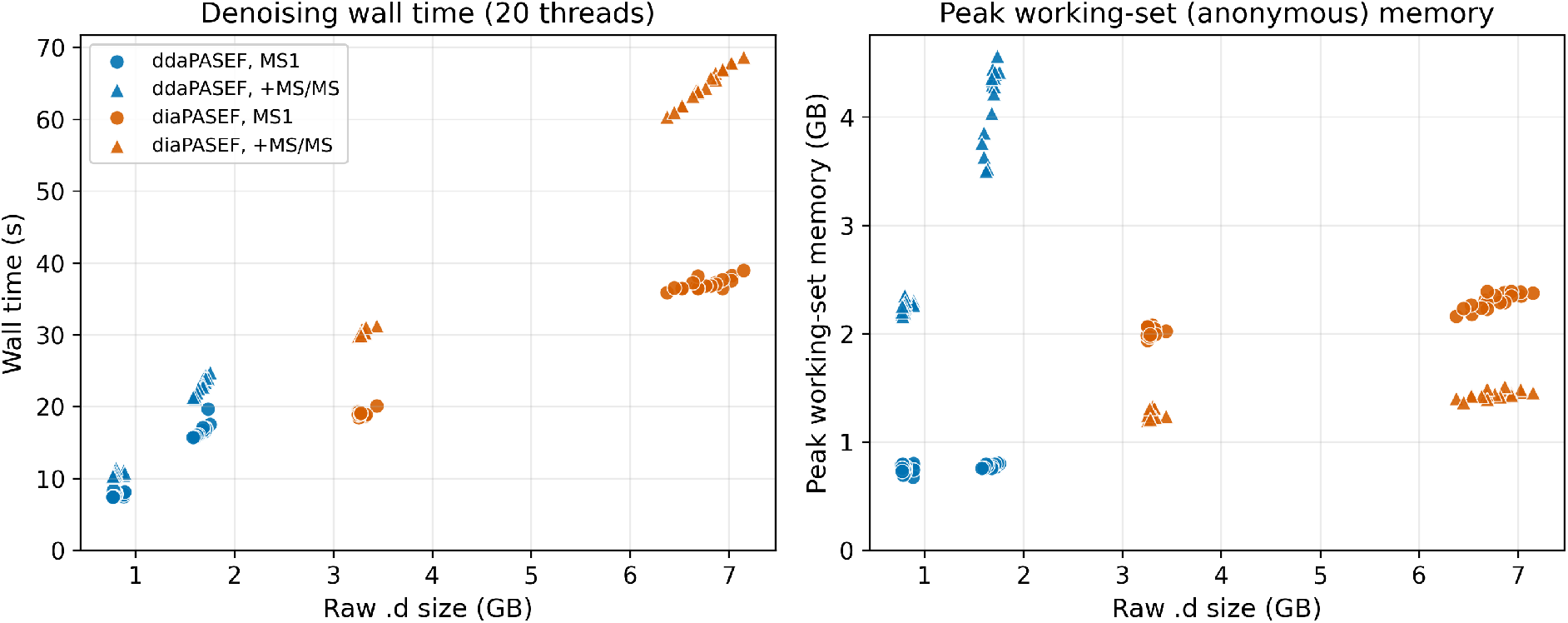
Runtime and memory across all 72 benchmark runs, both arms on one workstation (i7-12700H, 20 threads). Peak working-set memory is anonymous resident memory and excludes the memory-mapped input pages. Complete values are in Table S16.

The largest observed working set was 4.56 GB, for a 15-minute ddaPASEF file with MS/MS filtering. The working set remained below 2.4 GB even for the largest diaPASEF files, which were approximately 7 GB each. Total resident set size (RSS) reached 8.8 GB because it also includes file-backed pages from the memory-mapped input, which the operating system can reclaim. Per-acquisition timing and memory values are given in Table S16.

The tool writes the reduced frames to a new, standard Bruker `.d` directory. Every denoised directory in this benchmark was processed by the same Sage and DIA-NN workflows as its unmodified original. The measured processing times and memory requirements permit dnoise to run immediately after acquisition, before data transfer, analysis, replication, or archival. Each subsequent copy then inherits the storage saving.

### 3.7 Limitations

The streak filter assumes that genuine ions are sampled over enough consecutive mobility scans to form a recoverable run. The default parameters may therefore require relaxation for sparse or low-input data, including single-cell experiments.

The replicated benchmark remains controlled and narrow: one timsTOF Ultra 2 instrument, one laboratory, one 50 ng three-species sample, two gradients, and two acquisition modes. The difference in reduction between DDA and DIA is also confounded by on-instrument MS1 denoising having been enabled only for the primary DDA acquisition. Validation across laboratories, sample loads, and sparse acquisitions remains necessary. Every filter parameter is exposed in the configuration, so the operating point can be adapted to other sample types, experiments, and instruments, but whether the defaults or any adapted setting preserve quantification in those regimes is an empirical question this benchmark does not answer. Finally, dnoise removes points rather than providing lossless compression. Original files should be retained whenever future analyses may require signal outside the assumptions tested here.

## 4 Conclusions

dnoise removes points directly from native Bruker `.d` data while preserving the results of the tested proteomics workflows. In its default MS1-only mode, it removed the large majority of MS1 points and reduced the frame binary by 35–53% across the tested acquisitions. LFQ accuracy was preserved in both acquisition modes. ddaPASEF identification counts were unchanged, and diaPASEF precursor and protein-group counts changed only slightly. The resulting `.d` directories remained compatible with existing downstream software, and every tested processing run completed in 69 seconds or less. Optional MS/MS denoising reduced the data further but sacrificed several percent of peptide and protein-group identifications, making MS1-only denoising the recommended default. These results show that a substantial fraction of timsTOF frame data can be removed before transfer or archival without materially changing the result of the proteomics workflows tested here.

## Associated Content

### Data Availability Statement

The primary raw mass spectrometry data are publicly available from the ProteomeXchange Consortium via the PRIDE partner repository under accession PXD070049 (timsTOF Ultra 2, released 3 February 2026 and publicly accessible, with no reviewer credentials required), and resolve at ebi.ac.uk/pride/archive/projects/PXD070049. dnoise is open source (MIT licensed) at github.com/pgarrett-scripps/dnoise and published on crates.io. The exact software release used for the reported results is dnoise v0.1.0, archived at doi.org/10.5281/zenodo.21959649. The release tag and Zenodo record are the immutable public software references for this study. The complete dnoise parameters (Table S1), Sage configuration, and DIA-NN commands (Section S7) are given in the manuscript and Supporting Information. Together with the public raw data and software release, these materials permit the denoising and database-search stages to be repeated.

### Supporting Information

Default parameters, complete ddaPASEF and diaPASEF benchmark results, full label-free quantification accuracy tables, representative MS/MS frames and parameter sweeps, matched intensity-threshold controls, optional MS1 centroiding, plus performance, memory, software, hardware, configuration, and reproduction information (PDF).

## Author Information

### Author Contributions

P.T.G. conceived and implemented dnoise, designed and performed the benchmark experiments, analyzed the data, and drafted the manuscript. J.K.D. contributed mass spectrometry expertise and advised on the benchmark design and data interpretation. J.R.Y. supervised the project and revised the manuscript. All authors reviewed and approved the final version.

### Notes

The authors declare no competing financial interest. This work is a computational analysis of a publicly deposited commercial multi-species digest standard (PXD070049) and involved no human subjects or animal procedures.

## Acknowledgment

We thank Claire Delahunty, Ph.D., for a careful reading of the manuscript. This work was supported by the U.S. National Institutes of Health (grants R01 HL165168, R01 AG077046, R01 MH100175, and R01 AG075862 to J.R.Y., R01 MH132570 to L. Ye, and U01 AG088679 to S. A. Lipton) and by the Skaggs Graduate School of Chemical and Biological Sciences at The Scripps Research Institute (P.T.G.). During the preparation of this work the authors used Anthropic Claude large language models via the Claude Code interface for manuscript drafting and editing and for software-development assistance, including development of the analysis and figure-generation scripts. No AI tool directly generated an image or produced or altered experimental data: every figure and reported value was computed by those scripts from the acquired raw data. The authors reviewed and edited all content and take full responsibility for the content of the publication.

