## Supplemental Information for "dnoise: Fast Native Data Reduction for Bruker timsTOF"

The Supporting Information provides the default parameters, complete benchmark results, the matched intensity-threshold control, the optional MS1 centroiders, performance details, and information needed to reproduce the work. The three benchmark arms are original, MS1-only denoising, and MS1+MS/MS denoising. Unless stated otherwise, identifications were controlled at 1% FDR and quantification follows Section 2.7.

#### S1 Default parameters

| Parameter | Default | Meaning |
|---|---|---|
| `mz_half_width` | 3 | TOF-index window half-width |
| `min_feature_length` | 5 | minimum occupied scans in a kept MS1 run |
| `max_internal_gap` | 2 | empty scans bridged within an MS1 run |
| `iterations` | 2 | MS1 filtering passes |
| `halo_peak_fraction` | 0.15 | minimum fraction of local off-column maximum |
| `halo_mz_idx_half_width` | 80 | halo-box TOF-index half-width |
| `halo_scan_half_width` | 2 | halo-box scan half-width |
| `msms_mz_half_width` | 3 | MS/MS TOF-index window half-width |
| `msms_min_feature_length` | 3 | minimum occupied scans in an MS/MS run |
| `msms_max_internal_gap` | 8 | empty scans bridged in an MS/MS run |
| `msms_iterations` | 1 | MS/MS filtering passes |
| `ms1_polygon_mz_pad` | 5.0 | selection-polygon padding (Da) |
| `ms1_polygon_im_pad` | 0.05 | selection-polygon padding ($1/K_0$) |
| `dia_ms1_mz_pad` | 5.0 | diaPASEF MS1-window padding (Da) |
| `dia_ms1_im_pad` | 0.05 | diaPASEF MS1-window padding ($1/K_0$) |
| `smooth_mz_idx_half_width` | 2 | pre-watershed smoothing box (TOF index) |
| `smooth_scan_half_width` | 3 | pre-watershed smoothing box (scans) |
| `watershed_box_scan` | 10 | watershed: neighbor reach (scans) |
| `watershed_box_mz_idx` | 3 | watershed: neighbor reach (TOF index) |
| `watershed_max_tof_offset` | 10 | watershed: max follower distance from seed (TOF index) |
| `box_centroid_mz_idx_half` | 2 | box centroider: box half-width (TOF index) |
| `box_centroid_scan_half` | 2 | box centroider: box half-width (scans) |

Table S1: Principal dnoise parameters and compiled defaults used for the benchmark. Intensity floors were zero. The streak, halo, and acquisition-aware MS1 gates were enabled. MS/MS denoising was disabled in the default arm and enabled only in the MS1+MS/MS arm. The optional centroiders (`smooth_*`, `watershed_*`, `box_centroid_*`) are off by default and were enabled only for the comparison in Section S6. The complete default configuration is compiled into dnoise.

The selection sweep is described in the main text. Each setting re-denoised the six replicates from raw and carried them through the full Sage LFQ pipeline. Per-setting results are in Table S2.

| Setting | MS1 removed | Peptides ($\Delta$ raw) | Proteins (≥2 pep.) | Median CV | Med. $\lvert \log_2 \rvert$ | Within $\pm$10% |
|---|---|---|---|---|---|---|
| original (raw) | — | 25,340 | 3,412 | 0.082 | — | — |
| gap 1, len 3 | 72% | 25,806 (+466) | 3,487 | 0.082 | 0.015 | 88% |
| gap 1, len 4 | 80% | 25,807 (+467) | 3,485 | 0.083 | 0.019 | 86% |
| gap 1, len 5 | 85% | 25,558 (+218) | 3,452 | 0.083 | 0.023 | 84% |
| gap 1, len 6 | 88% | 25,706 (+366) | 3,456 | 0.084 | 0.027 | 82% |
| gap 1, len 7 | 89% | 25,377 (+37) | 3,422 | 0.085 | 0.031 | 80% |
| gap 2, len 3 | 64% | 25,755 (+415) | 3,474 | 0.081 | 0.011 | 89% |
| gap 2, len 4 | 72% | 25,809 (+469) | 3,479 | 0.081 | 0.013 | 89% |
| **gap 2, len 5** [a] | 78% | 25,745 (+405) | 3,482 | 0.081 | 0.015 | 88% |
| gap 2, len 6 | 81% | 25,774 (+434) | 3,480 | 0.081 | 0.017 | 87% |
| gap 2, len 7 | 84% | 25,465 (+125) | 3,445 | 0.081 | 0.018 | 86% |
| gap 3, len 3 | 57% | 25,389 (+49) | 3,446 | 0.080 | 0.009 | 89% |
| gap 3, len 4 | 66% | 25,605 (+265) | 3,470 | 0.081 | 0.010 | 89% |
| gap 3, len 5 | 71% | 25,731 (+391) | 3,486 | 0.081 | 0.011 | 89% |
| gap 3, len 6 | 75% | 25,931 (+591) | 3,505 | 0.081 | 0.013 | 88% |
| gap 3, len 7 | 78% | 25,659 (+319) | 3,482 | 0.081 | 0.013 | 88% |
| gap 3, len 9 | 82% | 25,608 (+268) | 3,474 | 0.081 | 0.015 | 87% |
| gap 3, len 11 | 85% | 25,557 (+217) | 3,438 | 0.081 | 0.017 | 85% |
| intensity T=41 | 46% | 25,320 (−20) | 3,447 | 0.080 | 0.006 | 91% |
| intensity T=54 | 61% | 25,401 (+61) | 3,454 | 0.081 | 0.010 | 91% |
| intensity T=72 | 75% | 25,523 (+183) | 3,458 | 0.082 | 0.020 | 89% |
| intensity T=104 | 87% | 26,008 (+668) | 3,485 | 0.086 | 0.049 | 79% |

Table S2: **Default-parameter sweep** (Condition A, 15-minute ddaPASEF, 6 replicates, 1% FDR). Each row reports MS1 points removed by the streak filter, quantified peptides (change vs. raw), quantified proteins (≥2 peptides), median peptide CV, and intensity fidelity to raw (median-aligned median $\lvert \log_2 \rvert$ and fraction within $\pm$10%). Identifications are identical across all settings because only MS1 is filtered. The adopted default is gap 2, length 5[a]. The intensity rows are per-point intensity thresholds at four removal levels, run as a control.

[a] Adopted default (`FilterParams::default`).

#### S2 Benchmark results

| Condition (% w/w) | Human | Yeast | *E. coli* |
|---|---|---|---|
| A | 65 | 30 | 5 |
| B | 65 | 15 | 20 |
| C | 65 | 3 | 32 |

Table S3: Three-species hybrid benchmark composition. Each condition was acquired in six replicates at each gradient and acquisition mode. The three pairwise comparisons span expected $\log_2$ ratios from −2.7 to +3.3.

| Stage | ddaPASEF 5 min | ddaPASEF 15 min | diaPASEF 5 min | diaPASEF 15 min |
|---|---|---|---|---|
| *MS1 points kept (% of raw)* | | | | |
| Acquisition gate only | 62.4% | 67.3% | 36.5% | 40.7% |
| Streak filter only | 36.5% | 38.2% | 17.3% | 19.9% |
| Gate + streak | 20.3% | 23.6% | 7.0% | 9.6% |
| Gate + streak + halo (default) | 19.7% | 22.8% | 6.7% | 9.2% |
| *Frame binary (% of raw)* | | | | |
| Acquisition gate only | 75.7% | 79.7% | 73.1% | 78.3% |
| Streak filter only | 59.3% | 62.7% | 65.7% | 71.7% |
| Gate + streak | 47.9% | 52.8% | 61.0% | 67.7% |
| Gate + streak + halo (default) | 47.5% | 52.3% | 60.9% | 67.5% |

Table S4: **Per-stage ablation of the default MS1 pipeline.** MS1 point retention and frame-binary size for one run (Condition A, replicate 1) per acquisition. All arms used the benchmark configuration. Stages were isolated by disabling the others (`--iterations 0` skips the streak filter, `--no-halo` the halo filter, and `--no-ms1-polygon`/`--no-dia-ms1-window` the gate). The gate is the ddaPASEF selection polygon or the padded diaPASEF isolation windows. MS/MS frames are untouched in every arm, which bounds how far the binary can shrink.

| Metric | 5-minute | | | 15-minute | | |
|---|---|---|---|---|---|---|
| | Orig. | MS1 | +MS/MS | Orig. | MS1 | +MS/MS |
| PSMs (1% FDR) | 195,701 | 195,701 | 172,947 | 680,963 | 680,963 | 636,123 |
| Identified peptides (1% FDR) | 17,814 | 17,814 | 15,624 | 45,287 | 45,287 | 42,249 |
| Identified protein groups (1% FDR) | 4,872 | 4,872 | 4,289 | 7,726 | 7,726 | 7,275 |
| Quantified proteins (≥2 peptides) | 1,633 | 1,639 | 1,725 | 3,962 | 4,108 | 3,985 |
| Quantified peptides | 7,656 | 7,557 | 8,454 | 28,467 | 29,555 | 28,404 |
| Median protein-level LFQ CV | 0.287 | 0.289 | 0.226 | 0.046 | 0.046 | 0.050 |
| $\log_2\left(\frac{A}{B}\right)$ human (exp. 0) | −0.15 | −0.16 | −0.12 | −0.13 | −0.13 | −0.13 |
| $\log_2\left(\frac{A}{B}\right)$ yeast (exp. +1) | +0.79 | +0.78 | +0.81 | +0.81 | +0.82 | +0.83 |
| $\log_2\left(\frac{A}{B}\right)$ *E. coli* (exp. −2) | −1.31 | −1.47 | −1.61 | −1.68 | −1.67 | −1.72 |

Table S5: **ddaPASEF identification and quantification summary.** Original, MS1-only, and MS1+MS/MS arms at both gradients (18 runs each). It contains the complete values summarized in Figures 2, 3 and 4.

| Metric | 5-minute | | | 15-minute | | |
|---|---|---|---|---|---|---|
| | Orig. | MS1 | +MS/MS | Orig. | MS1 | +MS/MS |
| Identified precursors (1% FDR) | 59,343 | 59,225 | 55,955 | 113,260 | 110,797 | 102,516 |
| Identified peptides (1% FDR) | 54,902 | 54,649 | 52,250 | 102,230 | 99,889 | 93,625 |
| Identified protein groups (1% FDR) | 9,537 | 9,516 | 9,155 | 11,717 | 11,533 | 10,953 |
| Quantified proteins (≥2 peptides) | 7,612 | 7,586 | 7,218 | 10,205 | 9,988 | 9,475 |
| yeast | 1,866 | 1,852 | 1,734 | 2,824 | 2,746 | 2,592 |
| *E. coli* | 605 | 608 | 538 | 942 | 899 | 823 |
| Quantified peptides | 52,696 | 52,457 | 49,912 | 100,346 | 97,883 | 91,638 |
| Median protein-level LFQ CV | 0.067 | 0.067 | 0.069 | 0.048 | 0.051 | 0.051 |
| $\log_2\left(\frac{A}{B}\right)$ human (exp. 0) | −0.12 | −0.12 | −0.13 | −0.12 | −0.12 | −0.12 |
| $\log_2\left(\frac{A}{B}\right)$ yeast (exp. +1) | +0.67 | +0.67 | +0.69 | +0.81 | +0.84 | +0.85 |
| $\log_2\left(\frac{A}{B}\right)$ *E. coli* (exp. −2) | −1.42 | −1.41 | −1.50 | −1.73 | −1.78 | −1.85 |

Table S6: **diaPASEF identification and quantification summary.** Counts are experiment-wide at 1% FDR from independent DIA-NN searches against the same denoise-independent predicted library. Expected human, yeast, and *E. coli* A/B $\log_2$ ratios are 0, +1, and −2.

| Acquisition | Metric | Gradient | Original mean | MS1 mean | Δ (MS1 − orig.) | 95% CI of Δ |
|---|---|---|---|---|---|---|
| ddaPASEF | Quantified proteins/ run | 5 min | 1629 | 1635 | +6.1 | [+4.4, +7.6] |
| ddaPASEF | Quantified proteins/ run | 15 min | 3961 | 4107 | +146.8 | [+146.2, +147.5] |
| diaPASEF | Identified precursors/ run | 5 min | 48936 | 49061 | +124.6 | [+30.7, +223.4] |
| diaPASEF | Identified precursors/ run | 15 min | 96743 | 93680 | −3062.5 | [−3207.0, −2928.7] |
| diaPASEF | Identified protein groups/run | 5 min | 8609 | 8609 | +0.2 | [−12.0, +12.6] |
| diaPASEF | Identified protein groups/run | 15 min | 10844 | 10586 | −257.6 | [−276.7, −238.7] |

Table S7: **Per-run paired results for original versus MS1-only denoising.** Each row reports the mean over 18 runs and the paired change with its bootstrap 95% confidence interval. The DDA metric is quantified proteins, and the DIA metrics are identified precursors and identified protein groups.

### S3 Label-free quantification

Tables S8 and S9 provide every species and condition-pair median used to assess LFQ accuracy. Precision is reported in the summary tables above as the median within-condition protein CV.

| Pair, species | exp. | 5 min | | | 15 min | | |
|---|---|---|---|---|---|---|---|
| | | orig. | MS1 | +MS/MS | orig. | MS1 | +MS/MS |
| A/B Human | +0.00 | −0.15 | −0.16 | −0.12 | −0.13 | −0.13 | −0.13 |
| A/B Yeast | +1.00 | +0.79 | +0.78 | +0.81 | +0.81 | +0.82 | +0.83 |
| A/B Ecoli | −2.00 | −1.31 | −1.47 | −1.61 | −1.68 | −1.67 | −1.72 |
| A/C Human | +0.00 | −0.18 | −0.19 | −0.16 | −0.21 | −0.20 | −0.20 |
| A/C Yeast | +3.32 | +2.52 | +2.66 | +2.60 | +2.59 | +2.61 | +2.70 |
| A/C Ecoli | −2.68 | −2.21 | −2.29 | −2.31 | −2.46 | −2.46 | −2.50 |
| B/C Human | +0.00 | −0.04 | −0.04 | −0.04 | −0.08 | −0.08 | −0.08 |
| B/C Yeast | +2.32 | +1.77 | +1.80 | +1.83 | +1.79 | +1.79 | +1.90 |
| B/C Ecoli | −0.68 | −0.75 | −0.75 | −0.76 | −0.75 | −0.75 | −0.76 |

Table S8: **ddaPASEF LFQ accuracy.** Observed median $\log_2$ ratio for every condition pair, species, arm, and gradient.

| Pair, species | exp. | 5 min | | | 15 min | | |
|---|---|---|---|---|---|---|---|
| | | orig. | MS1 | +MS/MS | orig. | MS1 | +MS/MS |
| A/B Human | +0.00 | −0.12 | −0.12 | −0.13 | −0.12 | −0.12 | −0.12 |
| A/B Yeast | +1.00 | +0.67 | +0.67 | +0.69 | +0.81 | +0.84 | +0.85 |
| A/B Ecoli | −2.00 | −1.42 | −1.41 | −1.50 | −1.73 | −1.78 | −1.85 |
| A/C Human | +0.00 | −0.25 | −0.24 | −0.26 | −0.23 | −0.25 | −0.24 |
| A/C Yeast | +3.32 | +2.16 | +2.11 | +2.21 | +2.61 | +2.72 | +2.79 |
| A/C Ecoli | −2.68 | −2.14 | −2.10 | −2.25 | −2.51 | −2.59 | −2.66 |
| B/C Human | +0.00 | −0.13 | −0.13 | −0.14 | −0.12 | −0.13 | −0.13 |
| B/C Yeast | +2.32 | +1.47 | +1.43 | +1.54 | +1.78 | +1.88 | +1.96 |
| B/C Ecoli | −0.68 | −0.68 | −0.68 | −0.72 | −0.77 | −0.79 | −0.80 |

Table S9: **diaPASEF LFQ accuracy.** Observed median $\log_2$ ratio for every condition pair, species, arm, and gradient.

The paired tables below use the same proteins in the original and default MS1-denoised arms. They report the accuracy change directly with the bootstrap confidence intervals described in Section 2.7.

| Gradient | Pair | Species | $n$ | Median orig. | Median MS1 | $\Delta$ (MS1 − orig.) | 95% CI of $\Delta$ |
|---|---|---|---|---|---|---|---|
| 5 min | A/B | *human* | 1183 | −0.146 | −0.161 | −0.015 | [−0.028, −0.001] |
| 5 min | A/B | *yeast* | 256 | +0.783 | +0.784 | +0.000 | [−0.024, +0.021] |
| 5 min | A/B | *E. coli* | 131 | −1.351 | −1.475 | −0.123 | [−0.214, +0.006] |
| 5 min | A/C | *human* | 1183 | −0.182 | −0.199 | −0.018 | [−0.031, +0.001] |
| 5 min | A/C | *yeast* | 256 | +2.521 | +2.622 | +0.101 | [+0.005, +0.180] |
| 5 min | A/C | *E. coli* | 131 | −2.240 | −2.290 | −0.050 | [−0.140, +0.024] |
| 5 min | B/C | *human* | 1183 | −0.035 | −0.044 | −0.009 | [−0.017, +0.005] |
| 5 min | B/C | *yeast* | 256 | +1.771 | +1.764 | −0.006 | [−0.053, +0.068] |
| 5 min | B/C | *E. coli* | 131 | −0.750 | −0.753 | −0.002 | [−0.034, +0.052] |
| 15 min | A/B | *human* | 2842 | −0.130 | −0.130 | −0.000 | [−0.002, +0.002] |
| 15 min | A/B | *yeast* | 751 | +0.808 | +0.818 | +0.010 | [+0.004, +0.015] |
| 15 min | A/B | *E. coli* | 330 | −1.692 | −1.665 | +0.027 | [−0.018, +0.068] |
| 15 min | A/C | *human* | 2842 | −0.212 | −0.205 | +0.007 | [+0.004, +0.011] |
| 15 min | A/C | *yeast* | 750 | +2.591 | +2.600 | +0.008 | [−0.023, +0.035] |
| 15 min | A/C | *E. coli* | 330 | −2.463 | −2.456 | +0.007 | [−0.023, +0.066] |
| 15 min | B/C | *human* | 2842 | −0.086 | −0.078 | +0.007 | [+0.004, +0.010] |
| 15 min | B/C | *yeast* | 750 | +1.788 | +1.785 | −0.003 | [−0.033, +0.020] |
| 15 min | B/C | *E. coli* | 330 | −0.753 | −0.753 | −0.000 | [−0.010, +0.013] |

Table S10: **Paired ddaPASEF LFQ accuracy changes under default MS1 denoising.** Each row uses the shared contributing proteins and reports the change in the median $\log_2$ ratio with its percentile-bootstrap 95% confidence interval.

| Gradient | Pair | Species | $n$ | Median orig. | Median MS1 | $\Delta$ (MS1 − orig.) | 95% CI of $\Delta$ |
|---|---|---|---|---|---|---|---|
| 5 min | A/B | *human* | 4965 | −0.125 | −0.123 | +0.002 | [+0.000, +0.005] |
| 5 min | A/B | *yeast* | 1795 | +0.675 | +0.675 | +0.000 | [−0.006, +0.008] |
| 5 min | A/B | *E. coli* | 563 | −1.450 | −1.436 | +0.013 | [−0.013, +0.041] |
| 5 min | A/C | *human* | 4966 | −0.251 | −0.246 | +0.006 | [+0.002, +0.009] |
| 5 min | A/C | *yeast* | 1285 | +2.192 | +2.171 | −0.021 | [−0.049, −0.002] |
| 5 min | A/C | *E. coli* | 563 | −2.178 | −2.140 | +0.038 | [−0.001, +0.058] |
| 5 min | B/C | *human* | 4973 | −0.134 | −0.130 | +0.003 | [+0.001, +0.005] |
| 5 min | B/C | *yeast* | 1285 | +1.512 | +1.483 | −0.030 | [−0.050, −0.006] |
| 5 min | B/C | *E. coli* | 722 | −0.684 | −0.686 | −0.001 | [−0.010, +0.009] |
| 15 min | A/B | *human* | 6165 | −0.120 | −0.123 | −0.004 | [−0.006, −0.002] |
| 15 min | A/B | *yeast* | 2683 | +0.814 | +0.838 | +0.023 | [+0.018, +0.029] |
| 15 min | A/B | *E. coli* | 859 | −1.750 | −1.799 | −0.049 | [−0.062, −0.028] |
| 15 min | A/C | *human* | 6164 | −0.235 | −0.248 | −0.013 | [−0.016, −0.010] |
| 15 min | A/C | *yeast* | 1984 | +2.666 | +2.741 | +0.075 | [+0.052, +0.094] |
| 15 min | A/C | *E. coli* | 859 | −2.545 | −2.603 | −0.059 | [−0.086, −0.042] |
| 15 min | B/C | *human* | 6169 | −0.120 | −0.128 | −0.008 | [−0.010, −0.006] |
| 15 min | B/C | *yeast* | 1984 | +1.848 | +1.915 | +0.067 | [+0.045, +0.090] |
| 15 min | B/C | *E. coli* | 1070 | −0.772 | −0.794 | −0.022 | [−0.028, −0.016] |

Table S11: **Paired diaPASEF LFQ accuracy changes under default MS1 denoising.** Each row uses the shared contributing proteins and reports the change in the median $\log_2$ ratio with its percentile-bootstrap 95% confidence interval.

| Gradient | Comparison | $n$ shared | Median CV orig. | Median CV arm | Yeast Δ (A/B) [95% CI] | E. coli Δ (A/B) [95% CI] |
|---|---|---|---|---|---|---|
| 5 min | MS1 | 48,637 | 0.159 | 0.176 | +0.067 [+0.056, +0.076] | −0.206 [−0.262, −0.152] |
| 5 min | MS1+MS/MS | 45,136 | 0.156 | 0.162 | +0.134 [+0.121, +0.144] | −0.380 [−0.443, −0.327] |
| 15 min | MS1 | 94,019 | 0.116 | 0.128 | +0.124 [+0.117, +0.132] | −0.313 [−0.342, −0.280] |
| 15 min | MS1+MS/MS | 87,160 | 0.111 | 0.119 | +0.176 [+0.168, +0.183] | −0.385 [−0.420, −0.354] |

Table S12: **Direct precursor MS1-area validation in diaPASEF.** Values use DIA-NN's `Ms1.Normalised` field on precursors shared between each denoised arm and the original arm. The table reports median precursor CV and paired A/B ratio changes for the regulated species with percentile-bootstrap 95% confidence intervals.

### S4 Optional MS/MS denoising

Representative frames illustrate that the retained points follow coherent fragment-ion mobility structure (Figures S1 and S2). Parameter sweeps show the tunable reduction-versus-identification tradeoff (Figure S3).

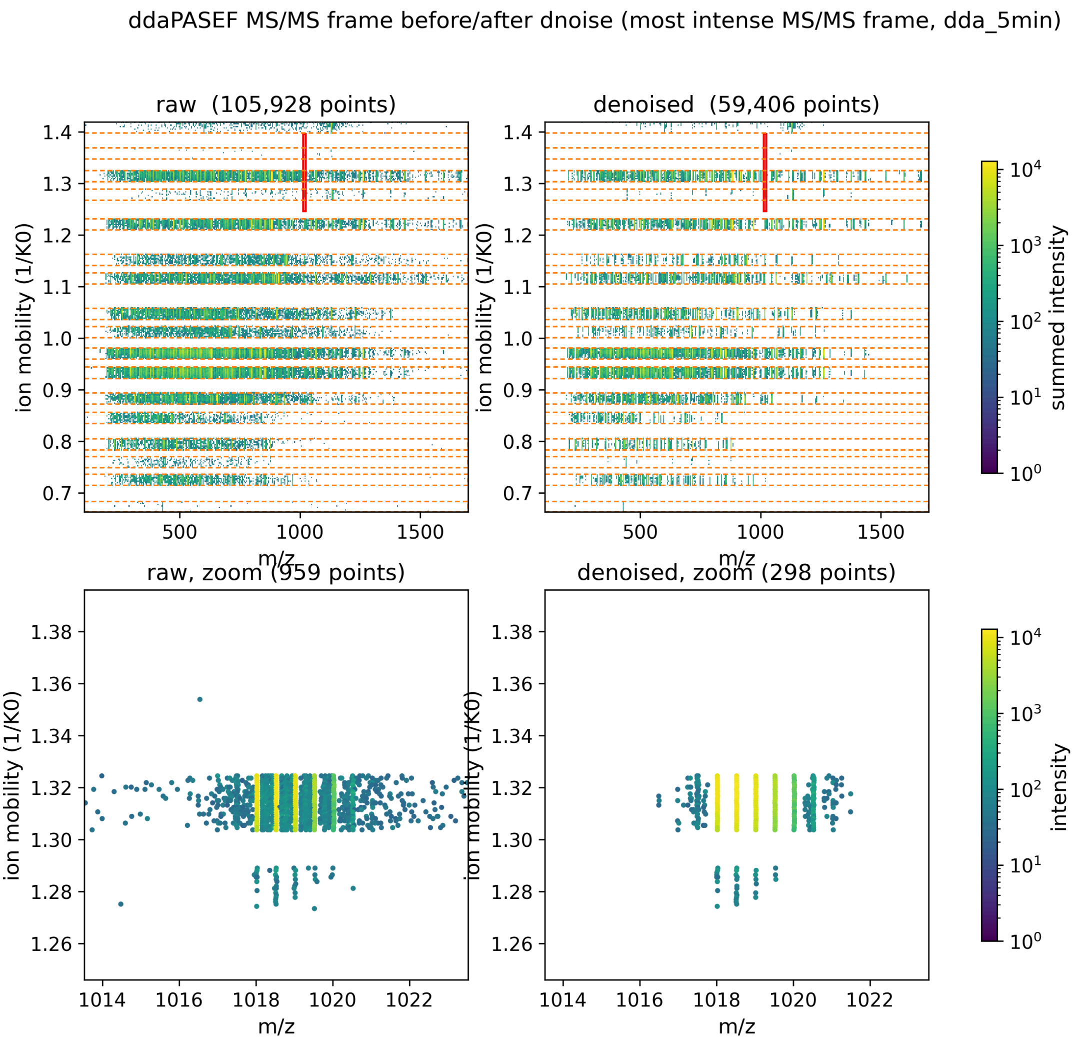


Figure S1: **Representative ddaPASEF MS/MS frame before and after denoising.** Dashed outlines show precursor-isolation events.

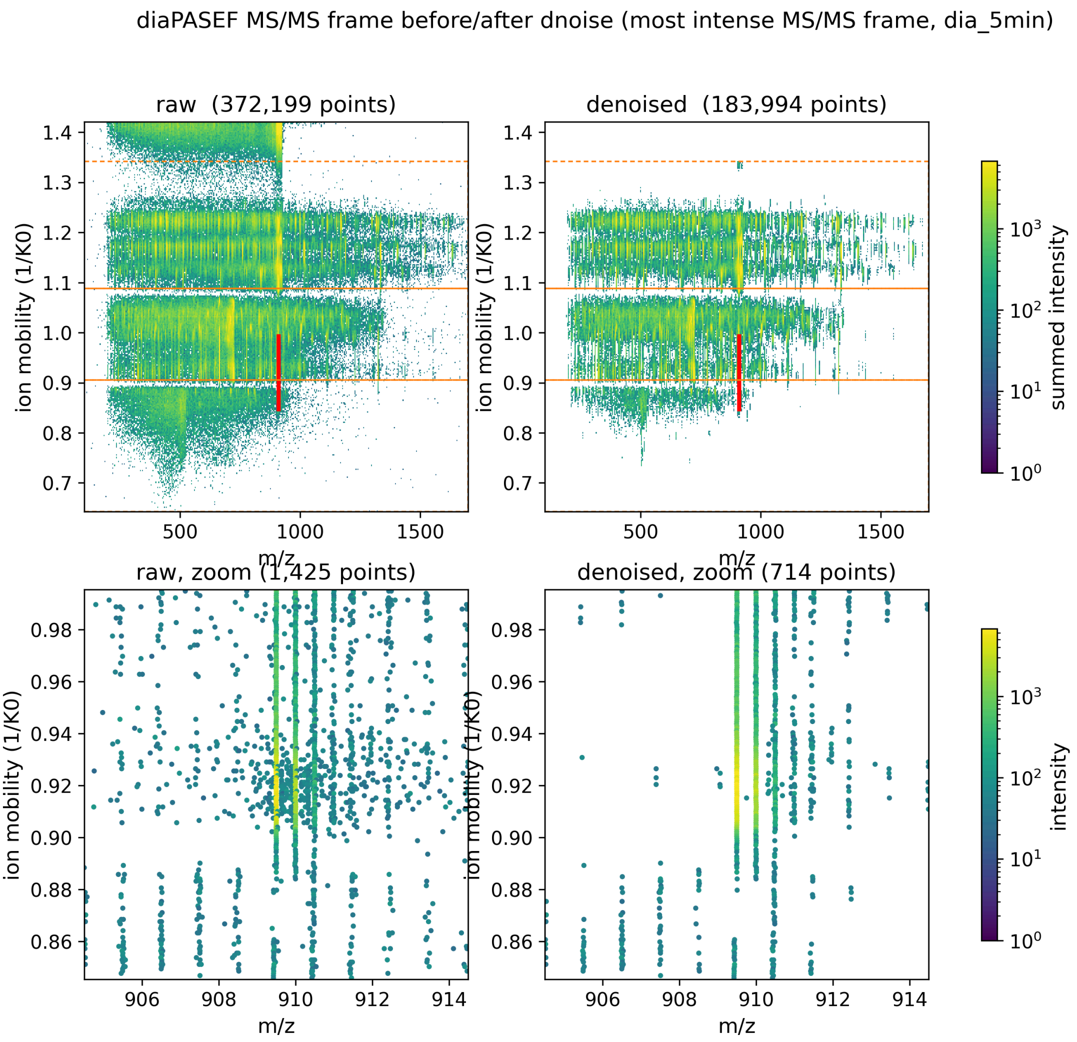


770 Figure S2: **Representative diaPASEF MS/MS frame before and after denoising.**
771 Dashed outlines show isolation windows. Identification counts across the complete benchmark
772 are reported in Tables S5 and S6.

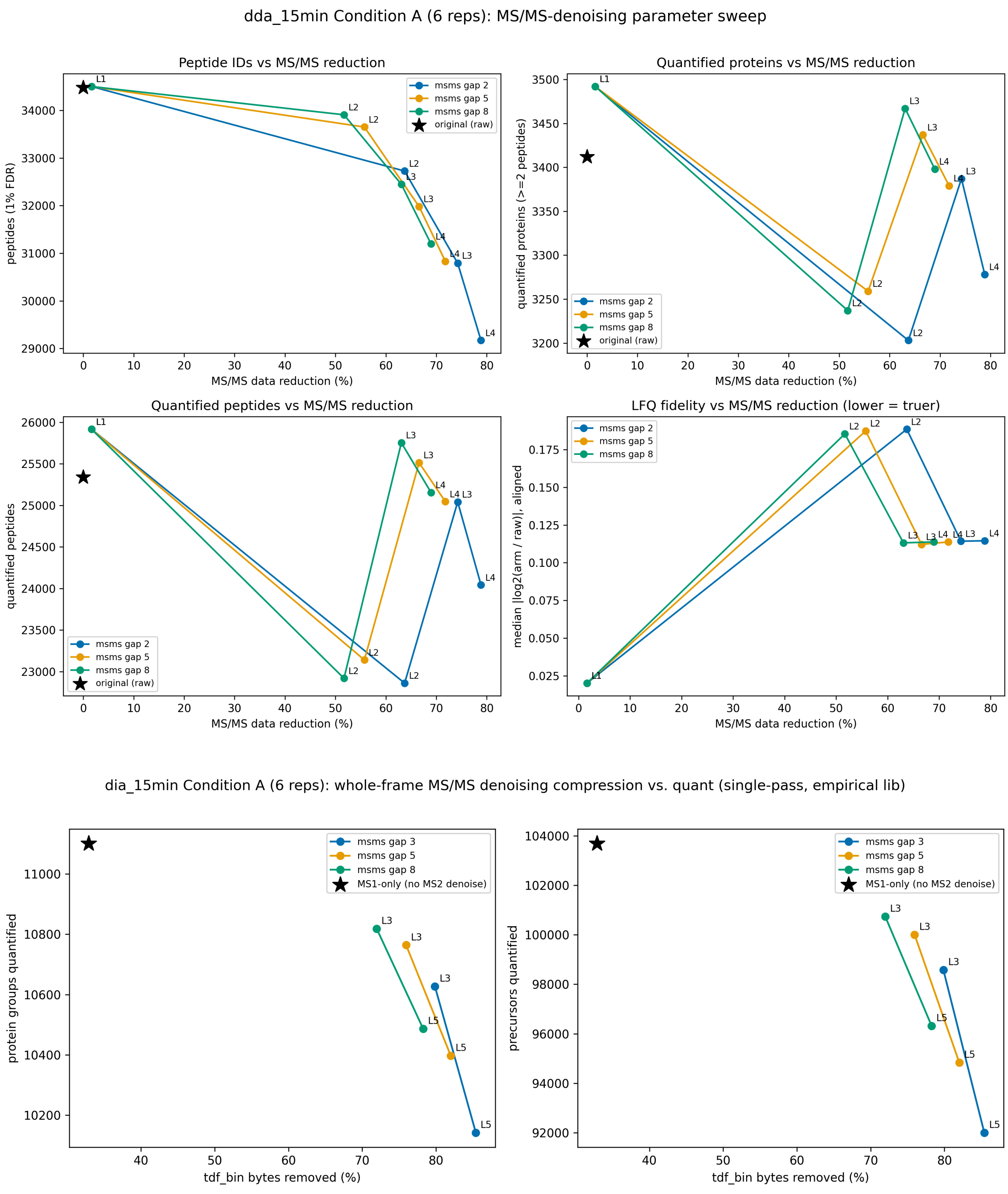


Figure S3: **MS/MS reduction-versus-identification tradeoff.** Top: ddaPASEF parameter sweep. Bottom: diaPASEF parameter sweep. The tested configuration was held fixed across the benchmark after selection as described in Section 2.2.

### S5 Matched intensity-threshold control

The control design is described in the main text: a strict per-point intensity threshold $T$, run through the same pipeline as the streak arms including the acquisition-aware gates, with $T$ calibrated separately for each acquisition to approximately match the streak arm's total point removal. The full numerical comparison is given in Table S13 and Table S14.

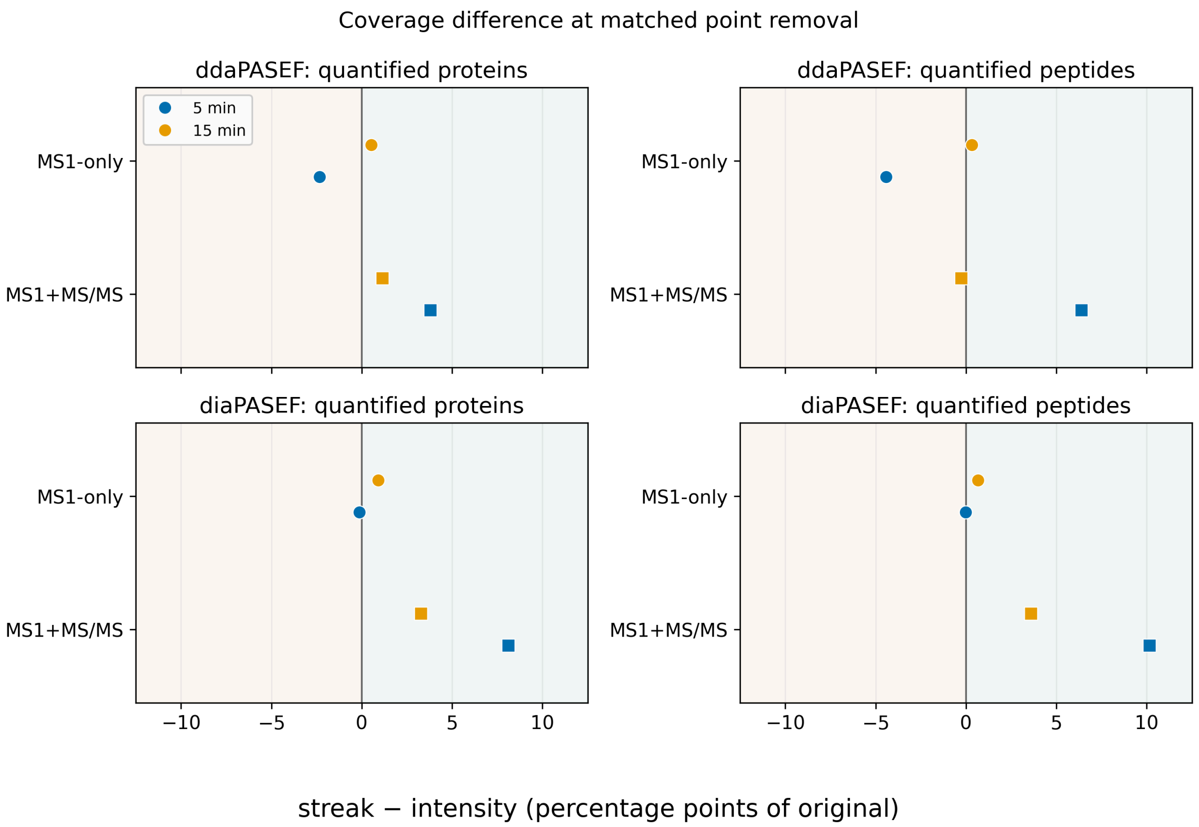


Figure S4: **Quantified-coverage difference between streak and matched intensity filtering.** Positive values favor the streak filter. The intensity threshold was calibrated separately for each acquisition to approximately match point removal.

To compare intensities directly, we retained peptide-run pairs with positive LFQ intensity in the original, default streak, and threshold arms, separately for the 5- and 15-minute ddaPASEF gradients. Original values defined equal-count abundance bins independently of both filters. At both gradients and across every bin, the streak filter preserved peptide LFQ intensity more closely than the threshold. Its advantage widened as abundance decreased, with the threshold showing its largest downward bias among the faintest peptides while the streak arm remained close to the original. In the faintest 15-minute decile the streak arm's median slightly exceeded the original (Figure S5), consistent with denoising removing interfering background from the extracted signal. This pattern is consistent with the streak filter retaining weak but mobility-coherent signal that falls below a fixed intensity cutoff. The streak and threshold arms removed 81.2% and 82.3% of MS1 points at 5 minutes, and 78.2% and 79.1% at 15 minutes. The absolute separation therefore includes the threshold arm's slightly greater point removal.

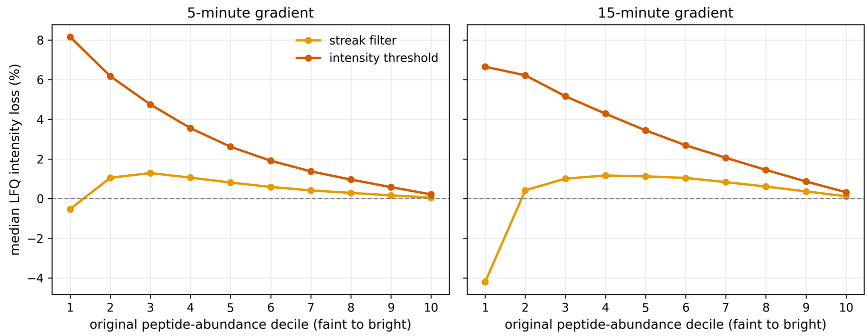


Figure S5: **Peptide LFQ intensity loss by abundance, streak filter versus matched intensity threshold.** Shared peptide-run pairs from all 18 runs at each ddaPASEF gradient were divided into equal-count deciles by original LFQ intensity. Points show the median percentage of original peptide intensity removed by each filter. The streak arm used the default settings in Table S1; the threshold arm used the deposited per-point thresholds, $T = 84$ at 5 minutes and $T = 79$ at 15 minutes. Both filters were restricted to MS1, so MS/MS spectra were unchanged. The control was approximately matched for point removal, not post hoc matched within each run.

Observed decoy attrition for the ddaPASEF searches is summarized in Figure S6. The DIA-NN report carries no decoys, so the diaPASEF equivalent is the expected false-positive count computed per arm from the reported posterior error probabilities. The accepted set is thresholded at 1% FDR, so that rate is pinned near 1% by construction in every arm, and the between-arm differences are changes in coverage (Table S6), not in error rate.

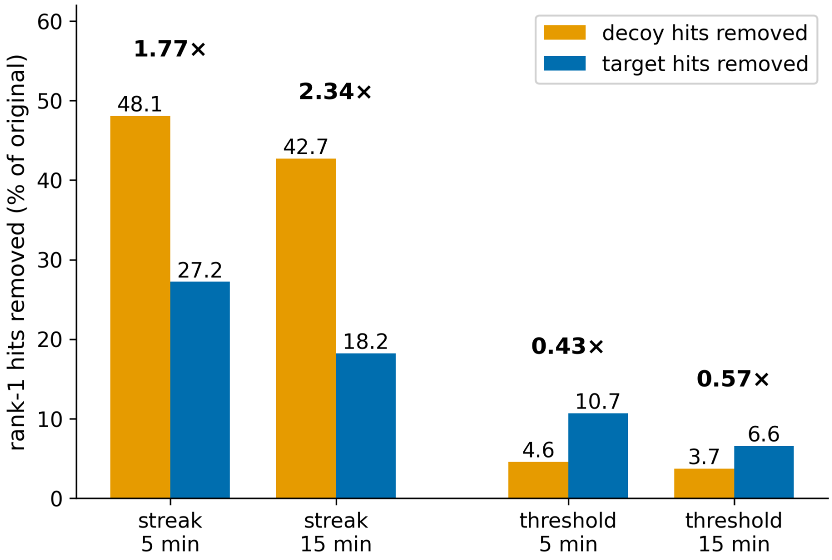


Figure S6: **Observed decoy versus target loss under fragment denoising, ddaPASEF.** A rank-1 decoy hit is a match to a sequence known to be absent from the sample, so decoy attrition counts false positives directly, with no score cutoff. The annotated ratio is decoy loss divided by target loss: above 1 a filter removes unidentifiable spectra faster than real evidence, below 1 it discards evidence faster. The MS1-only arms reproduce the original search exactly and are omitted.

| Metric | 5-minute | | | 15-minute | | |
|---|---|---|---|---|---|---|
| | Orig. | Streak | Intens. | Orig. | Streak | Intens. |
| **MS1-only** | | | | | | |
| Quantified proteins (≥2 peptides) | 1,633 | 1,639 | 1,677 | 3,962 | 4,108 | 4,087 |
| Quantified peptides | 7,656 | 7,557 | 7,896 | 28,467 | 29,555 | 29,463 |
| Median protein-level LFQ CV | 0.287 | 0.289 | 0.290 | 0.046 | 0.046 | 0.047 |
| $\log_2\left(\frac{A}{B}\right)$ *E. coli* (exp. −2) | −1.31 | −1.47 | −1.46 | −1.68 | −1.67 | −1.72 |
| **MS1+MS/MS** | | | | | | |
| Quantified proteins (≥2 peptides) | 1,633 | 1,725 | 1,663 | 3,962 | 3,985 | 3,940 |
| Quantified peptides | 7,656 | 8,454 | 7,965 | 28,467 | 28,404 | 28,478 |
| Median protein-level LFQ CV | 0.287 | 0.226 | 0.314 | 0.046 | 0.050 | 0.046 |
| $\log_2\left(\frac{A}{B}\right)$ *E. coli* (exp. −2) | −1.31 | −1.61 | −1.49 | −1.68 | −1.72 | −1.77 |

Table S13: **Streak filter versus matched intensity threshold in ddaPASEF.** The top block is MS1-only and the bottom block MS1+MS/MS. Thresholds were recalibrated at each gradient to approximately match point removal.

| Metric | 5-minute | | | 15-minute | | |
|---|---|---|---|---|---|---|
| | Orig. | Streak | Intens. | Orig. | Streak | Intens. |
| **MS1-only** | | | | | | |
| Quantified proteins (≥2 peptides) | 7,612 | 7,586 | 7,596 | 10,205 | 9,988 | 9,896 |
| Quantified peptides | 52,696 | 52,457 | 52,462 | 100,346 | 97,883 | 97,209 |
| Median protein-level LFQ CV | 0.0666 | 0.0673 | 0.0666 | 0.0477 | 0.0513 | 0.0499 |
| $\log_2\left(\frac{A}{B}\right)$ *E. coli* (exp. −2) | −1.42 | −1.41 | −1.39 | −1.73 | −1.78 | −1.79 |
| **MS1+MS/MS** | | | | | | |
| Quantified proteins (≥2 peptides) | 7,612 | 7,218 | 6,601 | 10,205 | 9,475 | 9,142 |
| Quantified peptides | 52,696 | 49,912 | 44,570 | 100,346 | 91,638 | 88,037 |
| Median protein-level LFQ CV | 0.0666 | 0.0688 | 0.0758 | 0.0477 | 0.0511 | 0.0490 |
| $\log_2\left(\frac{A}{B}\right)$ *E. coli* (exp. −2) | −1.42 | −1.50 | −1.68 | −1.73 | −1.85 | −1.96 |

Table S14: **Streak filter versus matched intensity threshold in diaPASEF.** The top block is MS1-only and the bottom block MS1+MS/MS. Searches used the same predicted library as the diaPASEF benchmark analysis.

### S6 Optional MS1 centroiding (watershed and box)

The two optional, mutually exclusive MS1 centroiders are characterized in full in Table S15 and Figure S7 (5-minute ddaPASEF). Their parameters and defaults are listed in Table S1. The pre-watershed box smoother (`smooth_*`) stabilizes seed order so that a noise-driven intensity spike does not split one ion into several centroids.

| Metric | Orig. | Watershed | Box |
|---|---|---|---|
| MS1 peaks removed | — | 98.2% | 90.5% |
| Frame binary (% of raw) | 100.0% | 33.9% | 40.3% |
| PSMs (1% FDR) | 195,701 | 195,701 | 195,701 |
| Identified peptides (1% FDR) | 17,814 | 17,814 | 17,814 |
| Identified protein groups (1% FDR) | 4,872 | 4,872 | 4,872 |
| Quantified proteins (≥2 peptides) | 1,633 | 1,612 | 1,713 |
| yeast | 270 | 264 | 290 |
| *E. coli* | 136 | 136 | 138 |
| Quantified peptides | 7,656 | 7,486 | 8,156 |
| Median protein-level LFQ CV | 0.287 | 0.273 | 0.280 |
| $\log_2(\frac{A}{B})$ human (exp. 0) | −0.15 | −0.15 | −0.15 |
| $\log_2(\frac{A}{B})$ yeast (exp. +1) | +0.79 | +0.79 | +0.81 |
| $\log_2(\frac{A}{B})$ *E. coli* (exp. −2) | −1.31 | −1.38 | −1.39 |

Table S15: **Optional MS1 centroiding** (5-minute ddaPASEF, 18 runs, 1% FDR). Original vs. the watershed and box centroiders. Both run on MS1 only, so PSM/peptide/protein-group identifications are identical to original. Quantification uses the two-peptides-per-protein rule. The frame point clouds are shown in Figure S7.

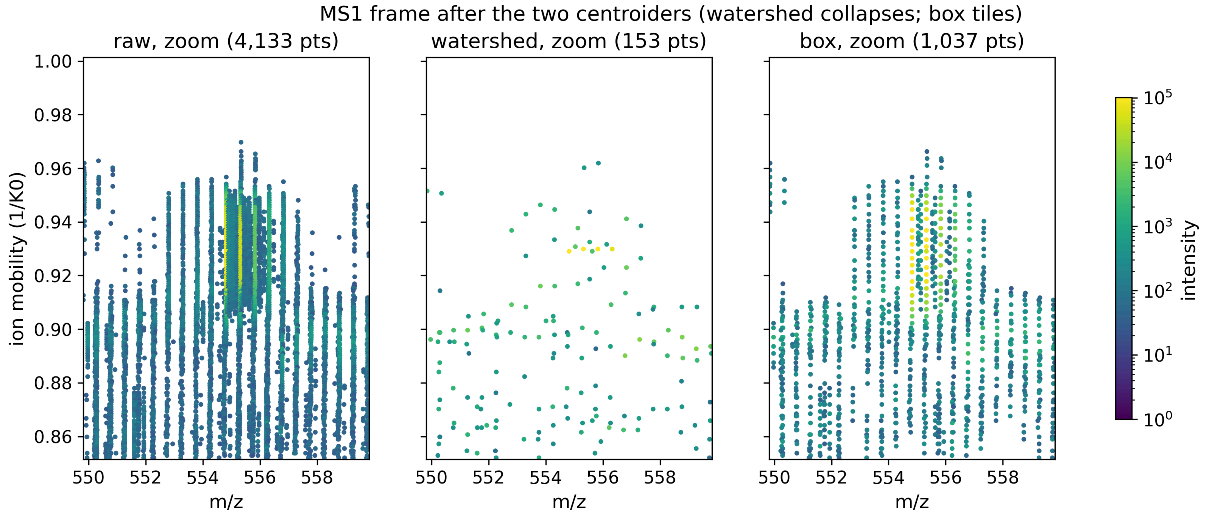


Figure S7: Optional MS1 centroiding (5-minute ddaPASEF, MS1-only, MS/MS untouched). Zoomed detail (10 m/z × 0.15 (1/K0)) of the most-intense MS1 frame: raw, then the watershed centroider (one centroid per ion streak) and the box centroider (several per streak, preserving the mobility profile). Metrics for both settings are given in Table S15.

### S7 Performance and reproducibility

| Acquisition | Arm | $n$ | Raw .d (GB) | Wall (s) | Total RSS (GB) | Working set (GB) | Input mmap (GB) |
|---|---|---|---|---|---|---|---|
| ddaPASEF, 5 min | MS1 | 18 | 0.80 (0.76–0.89) | 7.8 (7.4–8.3) | 1.37 (1.31–1.43) | 0.75 (0.68–0.80) | 0.74 (0.71–0.75) |
| ddaPASEF, 5 min | +MS/MS | 18 | 0.80 (0.76–0.89) | 10.7 (10.2–11.4) | 2.88 (2.76–2.97) | 2.26 (2.16–2.35) | 0.74 (0.71–0.75) |
| ddaPASEF, 15 min | MS1 | 18 | 1.69 (1.58–1.75) | 16.6 (15.7–19.6) | 2.21 (2.07–2.27) | 0.78 (0.76–0.80) | 1.60 (1.49–1.66) |
| ddaPASEF, 15 min | +MS/MS | 18 | 1.69 (1.58–1.75) | 23.6 (21.2–24.8) | 5.82 (4.98–6.09) | 4.32 (3.50–4.56) | 1.60 (1.49–1.66) |
| diaPASEF, 5 min | MS1 | 18 | 3.30 (3.25–3.44) | 18.8 (18.4–20.1) | 4.74 (4.57–4.86) | 2.03 (1.94–2.08) | 3.26 (3.15–3.38) |
| diaPASEF, 5 min | +MS/MS | 18 | 3.30 (3.25–3.44) | 30.3 (29.7–31.3) | 4.32 (4.17–4.53) | 1.25 (1.20–1.33) | 3.26 (3.21–3.40) |
| diaPASEF, 15 min | MS1 | 18 | 6.73 (6.38–7.15) | 36.8 (35.9–39.0) | 8.40 (7.89–8.78) | 2.30 (2.16–2.39) | 6.64 (6.27–6.94) |
| diaPASEF, 15 min | +MS/MS | 18 | 6.73 (6.38–7.15) | 64.2 (60.3–68.7) | 7.95 (7.53–8.39) | 1.43 (1.37–1.51) | 6.66 (6.31–7.08) |

Table S16: **Performance and memory per acquisition and arm.** Values are medians over 18 runs with ranges. Working set is peak anonymous resident memory. Total RSS also includes file-backed pages from the memory-mapped input, which the operating system can reclaim.

**Software and hardware.** dnoise 0.1.0 was built with Rust 1.97.1 and used `timsrust` 0.4.2 for frame I/O and `rayon` 1.12 for parallelism. ddaPASEF search and mobility-aware LFQ used Sage 0.15.0-beta.1. diaPASEF analysis used DIA-NN 2.2.0. Performance was measured on an Intel Core i7-12700H workstation (14 physical cores, 20 threads). The filters are deterministic integer operations, and builds made with Rust 1.91.0 and 1.97.1 produced byte-identical output.

**Configurations.** The complete default dnoise configuration is compiled into the tool (`FilterParams::default`) and listed in Table S1. Sage used the complete configuration below and reversed-sequence decoys with 1% FDR by the target-decoy strategy[25].

```
{
  "database": {
    "bucket_size": 8192,
    "enzyme": {
      "missed_cleavages": 2,
      "min_len": 7,
      "max_len": 30,
      "cleave_at": "KR",
      "restrict": "P",
      "c_terminal": true,
      "semi_enzymatic": false
    },
    "fragment_min_mz": 150.0,
```

```
    "fragment_max_mz": 2000.0,
    "peptide_min_mass": 500.0,
    "peptide_max_mass": 5000.0,
    "ion_kinds": ["b", "y"],
    "min_ion_index": 2,
    "static_mods": { "C": 57.0215 },
    "variable_mods": { "M": [15.9949] },
    "max_variable_mods": 2,
    "decoy_tag": "rev_",
    "generate_decoys": true,
    "fasta": "data/fasta/hybrid.fasta"
  },
  "precursor_tol": { "ppm": [-20.0, 20.0] },
  "fragment_tol": { "ppm": [-20.0, 20.0] },
  "isotope_errors": [-1, 2],
  "deisotope": true,
  "chimera": false,
  "wide_window": false,
  "predict_rt": true,
  "min_peaks": 5,
  "max_peaks": 150,
  "min_matched_peaks": 3,
  "max_fragment_charge": null,
  "report_psms": 1,
  "quant": {
    "lfq": true,
    "lfq_settings": {
      "peak_scoring": "Hybrid",
      "integration": "Sum",
      "spectral_angle": 0.7,
      "ppm_tolerance": 8.0,
      "mobility_pct_tolerance": 3.0,
      "combine_charge_states": true
    }
  },
  "output_directory": "results/<arm>"
}
```

The DIA-NN library was predicted once from the benchmark FASTA before denoising and then held fixed across all arms. The portable forms of the library-generation and per-arm search commands were:

```
diann-linux --fasta data/fasta/hybrid_diann.fasta --fasta-search \
  --gen-spec-lib --predictor --cut "K*,R*" --missed-cleavages 1 \
  --min-pep-len 7 --max-pep-len 30 --min-pr-charge 2 \
  --max-pr-charge 4 --min-pr-mz 300 --max-pr-mz 1800 \
  --min-fr-mz 200 --max-fr-mz 1800 --met-excision --unimod4 \
  --var-mods 1 --threads 20 --out-lib hybrid.tsv

diann-linux --f <each-arm.d> --lib hybrid.predicted.speclib \
  --fasta data/fasta/hybrid_diann.fasta --reanalyse --matrices \
  --qvalue 0.01 --threads 20 --out results/<arm>/report.parquet
```

**Availability.** Raw `.d` files and sample metadata are available from PRIDE PXD070049. dnoise is MIT licensed and available from `https://github.com/pgarrett-scripps/dnoise` and crates.io. Version 0.1.0 is the exact software release used here and is archived at `https://doi.org/`

10.5281/zenodo.21959649. The immutable public release tag is https://github.com/pgarrett-scripps/dnoise/releases/tag/v0.1.0. The exact Sage configuration and DIA-NN commands are reproduced above so the denoising and database-search stages can be repeated from the public raw data.
